# Progressive Changes in Functional Connectivity between Thalamic Nuclei and Cortical Networks Across Learning

**DOI:** 10.1101/2024.08.26.609333

**Authors:** Chelsea Jarrett, Katharina Zwosta, Xiaoyu Wang, Uta Wolfensteller, Juan Eugenio Iglesias, Katharina von Kriegstein, Hannes Ruge

**Author notes:** Shared last authorship (Katharina von Kriegstein, Hannes Ruge).

## Abstract

The thalamus is connected to cerebral cortex and subcortical regions, serving as a node within cognitive networks. It is a heterogeneous structure formed of functionally distinct nuclei with unique connectivity patterns. However, their contributions to cognitive functioning within networks is poorly understood. Recent animal research suggests that thalamic nuclei such as the mediodorsal nucleus play critical roles in goal-directed behaviour. Our aim was to investigate how functional integration of thalamic nuclei within cortical and subcortical networks changes whilst transitioning from more controlled goal-directed behaviour towards more automatic or habitual behaviour in humans. We analysed functional magnetic resonance imaging (fMRI) data from a stimulus-response learning study to investigate functional connectivity (FC) changes across learning between thalamic nuclei with cortical networks and subcortical structures in healthy subjects. We defined subcortical regions-of-interest (ROIs) individually in native space, segmenting the thalamus into 47 nuclei and segmenting 38 subregions within the basal ganglia and hippocampus. Additionally, we defined 12 cerebral cortex ROIs via maximum-probability network templates. Learning-related connectivity changes were examined via ROI-to-ROI functional network analysis. Our results showed that learning was associated with: 1) decreasing FC between the frontoparietal network and higher order thalamic nuclei; 2) increasing FC between the cingulo-opercular network and pulvinar nuclei, 3) decreasing FC between the default mode network (DMN) and right mediodorsal nuclei; 4) increasing FC between the DMN and left mediodorsal nuclei; 5) altered functional connectivity between thalamic nuclei and putamen subregions, and 6) increasing intrathalamic FC. Together, this suggests that several thalamic nuclei are involved in the learning-related transition from controlled to more automatic behaviour.

## 1 Introduction

Learning can be accompanied by a transition from goal-directed toward automatic control of behaviour and ultimately in the development of habits (Ersche, Lim, Ward, Robbins, & Stochl, 2017; B. Gardner, 2015; Yamada & Toda, 2023). This transition is known to be accompanied by changes in cerebral cortex networks and subcortical regions such as the striatum (van der Straten, Van Leeuwen, Denys, Marle, & van Wingen, 2020). How the thalamus is involved in the transition is largely unknown. This is surprising, as the mammalian brain is a thalamocortical system, with every cerebral cortex region receiving afferents from, and sending efferents back to, the thalamus (K. Hwang, Bertolero, Liu, & D’ Esposito, 2017). Additionally, the thalamus is extensively connected to the basal ganglia and other subcortical structures (Minagar et al., 2013) and is therefore in prime position to flexibly reconfigure large-scale neural networks (Kawabata et al., 2021; Saalmann & Kastner, 2015). As such, the thalamus might facilitate goal-directed behaviour through coordination of cognitive networks (Hwang *et* al. 2017) and cortico-striato-thalamo-cortical (CSTC) loops (Calzà et al., 2019) via its connections and cortical synchrony (Portoles et al., 2022), and might also contribute to automatic processes (Gremel & Lovinger, 2017).

Despite its function as network hub (Kawabata *et al*. 2021), thalamic roles in goal-directed behaviour within humans has been mostly ignored due to technological difficulties regarding small size and deep location (Calamante et al., 2013; Fama & Sullivan, 2015). Furthermore, most research investigates it as a whole structure (Goldstone et al., 2018) despite comprising around 60 cytoarchitectonically and functionally distinct nuclei with unique connectivity patterns with cortical and subcortical structures (Fama & Sullivan, 2015).

There is general consensus that higher order thalamic nuclei include mediodorsal (MD) and pulvinar subregions (Penner et al., 2018), alongside proposals to include anterior, lateral dorsal (Perry, Lomi, & Mitchell, 2021), ventral anterior, and ventral lateral nuclei (Sugiyama et al., 2018). MD nuclei are involved in executive functioning, decision-making and cognitive tasks (Pergola et al., 2018). They are critical for maintaining task representations and action-outcome contingencies (Fresno et al., 2019) and facilitate cognitive flexibility via initiating switching between cerebral cortex representations (Rikhye et al., 2018). This involves the frontoparietal network (FPN) (Li et al., 2022), but also default mode network (DMN) and salience networks (SN) (Niu et al., 2022). However, insight rests mostly on experiments in animals, not human subjects.

We investigated how integration of thalamic nuclei within networks might change across learning. Due to the prominent role of the MD nuclei within goal-directed behaviour and executive functioning, we expected these nuclei in particular to contribute to the transition from goal-directed toward automatic behaviour, potentially together with other higher thalamus regions. Furthermore, the MD nuclei are the largest thalamic nuclei in rodents, with even greater relative size within humans (Onishi, Kikuchi, Abe, Tokuhara, & Shimogori, 2022; Pergola et al., 2018). Mean volumes of the MD nuclei have been reported to be 1986 mm³ in healthy human participants (Danos et al., 2003), with such volumes thereby easily falling within the spatial resolution of standard human functional magnetic resonance imaging.

A recent fMRI study investigated transition from goal-directed to automatic behaviour across learning of stimulus-response associations by measuring changes in local BOLD (blood-oxygen dependent) responses and functional connectivity and tested whether learning-related changes predicted putative behavioural markers of habit strength (Zwosta et al., 2018). This revealed connectivity changes across cerebral cortex and basal ganglia during transition from more controlled to more automatic behaviour, alongside decreased thalamic responses and altered functional connectivity between thalamus and putamen. However, functional contributions of distinct thalamic nuclei underlying these alterations were not investigated not least due to a lack of analytical precision.

We therefore re-analysed data obtained within this study, and focused upon functional connectivity alterations between thalamic nuclei, cortical networks and subcortical regions across learning. In contrast to the original analysis approach, we now employed the methodological means to optimize analytical precision in order to obtain as much anatomical detail from standard functional images as possible.

## 2 Materials and Methods

### 2.1 Subjects

The sample originally used by Zwosta et al. (2018) consisted of a total of 53 subjects (29 female, 24 male) with a mean age of 23.5 years, and a range of 19-32 years. Here, we excluded one subject due to missing data in the nucleus accumbens region. All subjects were right-handed and had normal or corrected-to-normal vision.

The experimental protocol was approved by the Ethics Committee of the Technische Universität Dresden [EK306082011]. All subjects gave written informed consent prior to the experiment and were compensated 8€ per hour, in addition to any money earned throughout the experiment.

### 2.2 Experimental Procedure

The experimental paradigm was formed of three consecutive phases of which the second phase is of primary relevance for the present paper (Figure 1). In phase 1, goal-directed behaviour was established followed by phase 2 which was designed such that participants transitioned from initial goal-directed towards more automatic behaviour. In phase 3, goal-directed behaviour established in phase 1 was put into competition with more automatic behaviour established in phase 2.

**Figure 1.**
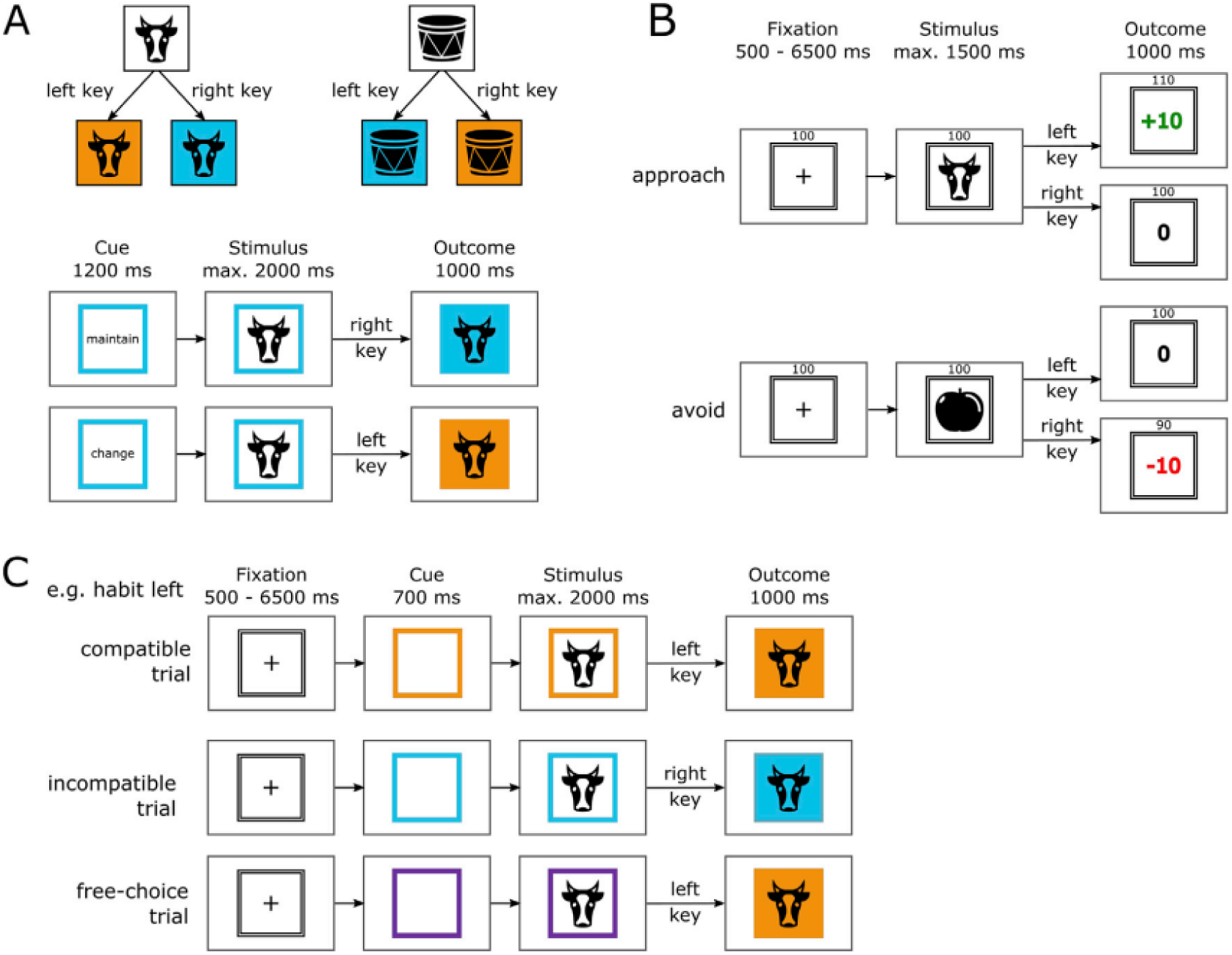
The experimental paradigm within Zwosta *et al*. (2018). The experiment consisted of three consecutive phases. In the present paper, we focused on fMRI data acquired during Phase 2. (A) Phase 1 established goal-directed behaviour directed towards different colour outcomes (outside the MRI machine). The panel shows examples of the instructed stimulus-response-outcome associations with two exemplary trials from Phase A. Subjects were instructed to learn the association between object categories (artificial, natural) and the respective colour and gave responses via button press. In a subsequent test they were asked to either maintain or change the colour association. (B) During Phase 2, novel stimulus-response mappings were learned for a subset of the stimuli used in Phase 1. Learning occurred via approach or avoidance and practiced inside the MRI machine. The panel shows exemplary trials from Phase 2, which consisted of approach and avoid trials. (C) Phase 3 placed goal-directed responses from Phase 1 and habitual responses from Phase 2 in competition with each other in order to test habit strength. The panel shows examples for compatible, incompatible, and free-choice trials within Phase C.

Phase 1 took place outside of the MRI machine, and goal-directed behaviour was established through the learning of 10 hierarchical stimulus-response-outcome (S-R-O) associations involving 10 distinct stimuli, 2 distinct responses, and 2 distinct colour outcomes. Phase 2 then took place within the MRI machine and included 8 of the visual stimuli from Phase 1, but now required subjects to learn novel stimulus-response associations by trial and error and with reinforcement. Here, the crucial assumption was that behaviour would be goal-directed initially (i.e., given a particular stimulus, select the action which will yield a rewarding outcome or enable the successful avoidance of an aversive outcome) and that this behaviour would gradually transition toward more automatic behaviour (i.e., the action will be generated ‘reflexively’ in response to the stimulus with increasingly less attention paid to the outcome). Finally, Phase 3 placed both goal-directed behaviour established in Phase 1 and habits established in Phase 2 into competition with each other within incompatible trials (which had required different responses in Phase 1 than they did in Phase 2) in order to quantify the habit strength acquired in Phase 2.

The present paper primarily focused upon the second phase with the aim to investigate thalamic contributions to the transition from goal-directed behaviour to more automatic behaviour. Here, across learning and practice, subjects were supposed to gradually transition from more controlled or goal-directed behaviour towards ever more automatic behaviour through learning trials and successful repetitions.

#### Phase 1: Establishment of goal-directed behaviour, outside of the MRI machine

This phase aimed at establishing a representation of hierarchical (here: stimulus-dependent) response-outcome (R-O) associations as the foundation of goal-directed behaviour (de Wit, Ostlund, Balleine, & Dickinson, 2009; Griffiths, Morris, & Balleine, 2014), while at the same time ensuring that no stable stimulus-response (S-R) habits could be formed (in contrast to phase 2 where this was the case).

The task utilised ten visual stimuli which were assigned into two categories: “artificial” and “natural”. These consisted of black and white, vertically symmetrical icons of different objects. Artificial stimuli included a ball, car, computer mouse, cupboard, and scissors, whereas natural stimuli included a cow, lungs, mushroom, snowflake, and a tree. When each image was presented on a screen, subjects were requested to press a corresponding key on a keyboard, i.e. by using the left index finger to press the key “D” or using the right index finger to press the key “K”. Responding to visual stimulus within one category with the right key would lead to a blue outcome colour whereas responding to the same category of visual stimulus with the left key would lead to an orange outcome colour. This R-O association was inverted for the other visual stimulus category, so that pressing the right key would lead to an orange outcome colour whereas pressing the left key would lead to a blue outcome colour. This inversion was designed to exclude habitualization processes, as for each stimulus, each of the two possible responses were equally often correct. Hence, the correct response was determined by the hierarchical relationship between stimulus category and intended goal (i.e. the anticipated colour outcome). Neither stimulus category (artificial or natural) alone nor intended goal alone, were sufficient to determine the correct response on a given trial.

Phase 1 began with subjects looking at an instruction screen which displayed all visual stimuli belonging to each group and the assigned R-O relationships. Next, the task was explained using text and illustrations. During this instruction phase, the experimenter stayed with each subject and during an additional 20 practice trials in order to answer any questions and ensure that the subject understood the instructions given.

Subsequently, subjects entered the testing phase. Here, they underwent a total of 240 trials, as outlined in Figure 1A. Each trial began with a cue on the screen, which was the German word for “change” or “maintain” surrounded by a square frame with either a blue or an orange colour. This colour would be the outcome colour achieved within the previous trial (or in the case of the first trial, would be a random colour). This cue would then be followed 1200 ms later by a visual stimulus from either the natural or artificial image categories. Here, the subject would utilise their knowledge of the hierarchical S-R-O associations learnt during the training phase in order to select the appropriate key in order to either ‘change’ or ‘maintain’ the colour as given by the coloured frame. The subjects had a response window of 2000 ms. Responses would cause the background of the stimulus to turn to the respective outcome colour for 1000 ms. Incorrect responses would generate the word ‘Error’ displayed below the square and the incorrect outcome colour would be shown. The subject would then have to immediately repeat the trial until they gave the correct response.

#### Phase 2: Stimulus-response learning, within the MRI machine

In this phase subjects were required to learn novel stimulus-response associations based on feedback in the form of either reward or punishment. Extensive practice of each S-R association was supposed to establish habitual approach or avoidance behaviour, respectively. Four artificial and four natural stimuli were re-used from Phase 1 (instead of the original 10 visual stimuli). At the beginning of phase 2, subjects were informed that the categories of the stimuli were no longer relevant and that during this phase they were required to discover and learn the correct key responses by trial and error. For both of the stimulus categories, each of the four stimuli belonging to each category was associated with one of the four combinations of correct responses (left or right key press) and outcome type (reward or punishment avoidance). Subjects obtained points by pressing the correct key response to four of the stimuli, whilst for the other four stimuli they would avoid losing points by pressing the correct key. Subjects were also informed that a faster response speed would also lead to gaining additional money to be obtained at the end of the experiment. Subjects received one cent per ten points and one cent per ms of their average response time being below 550 ms.

In Phase 2, each trial began with a fixation cross on the screen which was framed by a black-and-white lined square. This fixation cross was displayed for a variable duration, which ranged from 500 ms to 6500 ms (Figure 1B). A visual stimulus from one of the two image categories would then be presented, for a maximum of 1500 ms. The point outcome would be displayed immediately upon the response given by the subject. Rewards were 10+ points which were printed on the screen in green colour, whereas punishments consisted of -10 points which were printed in red. Outcomes of zero points were printed in black. Failure to respond during the response window would generate the unfavourable outcome, whereby in approach trials they would gain zero points (“0”) and in avoidance trials they would lose 10 points (“-10”). This outcome would remain on the screen for 1000 ms. Subjects were able to view their current number of points during the whole trial as this information was displayed at the top of the screen.

Phase 2 consisted of seven task blocks with 112 trials each (14 per stimulus). This formed a total of 784 trials, which was 98 per stimulus. The total number of points collected during each block would be displayed at the end of each block, and the mean response time for correct responses was also displayed at the end of each block.

#### Phase 3: Goal-directed vs habitual behaviour; testing habit strength within the MRI scanner

Both goal-directed behaviour from Phase 1 and habitual behaviour from Phase 2 were put into competition in order to assess inter-individual habit strength. Subjects were informed at the start of this phase that the rules of the prior phase 2 no longer applied, in that they could no longer gain or lose points. In this way, the contingency between stimulus, response and (monetary) outcome was removed, whereas the contingencies within Phase 1 were re-instructed. All ten stimuli from Phase 1 were used in Phase 3 (Figure 1C). A fixation cross was shown on the screen, which was followed by a coloured frame. This time, the frame was either orange or blue as within Phase A but could also be purple to indicate a free choice trial, whereby subjects could select either the left “D” or right “K” key. One of the ten visual stimuli then appeared on the screen. If the frame was blue or orange, then subjects were required to press the response which would lead to the outcome colour for the displayed stimulus in accordance with the R-O contingencies learnt within Phase 1 during goal-directed trials. A total of 6% of trials were catch trials, in order to prevent subjects selecting a response prior to the presentation of the stimulus. Subjects were also told not to decide how to respond before they were presented with the stimulus.

For all trials, the fixation cross was displayed for a variable ITI from 500 ms to 6500 ms and followed by a cue frame for 700 ms. The stimulus was displayed for up to 2000 ms or until the subject responded. After a response was provided, the outcome colour would be displayed for 1000 ms.

Phase 3 consisted of 384 trials in total. There was a total of eight different trial types of interest, each appearing in a random order. All trials required goal-directed responding as established in phase 1. This included ‘compatible’ trials for which the habitual response from phase 2 was identical to the required goal-directed response because it had been previously rewarded (approach compatible, 48 trials) or not punished (avoid compatible, 48 trials). There were also ‘incompatible’ trials for which the habitual response from phase 2 did not match the required goal-directed response and thereby caused a conflict between goal-directed response and the trained habitual response (approach incompatible or avoid incompatible, both 48 trials). Lastly, there were trials for which no habitual response existed as the stimulus had not been used within Phase 2 (no training, goal only, 48 trials). Additionally, there were free choice trials. For one type of free choice trials, the stimulus for which a habitual response was trained during Phase 2 was displayed (approach free-choice or avoid free-choice, each 48 trials). For another type of free-choice trials, the stimulus had not been used in the Phase 2 (no training free-choice, 24 trials). The difference between response times in incompatible vs compatible trials was used as a potential index of habit strength during the connectivity analysis. We call this difference between incompatible vs compatible trials the compatibility effect. The data from no-habit trials and the free choice trials were not used any further.

### 2.3 MRI Data Acquisition

MRI data was acquired on a Siemens 3T whole body Trio System (Erlangen, Germany) with a 32 channel head coil. Ear plugs were utilised in order to minimise MRI scanner noise. After the experimental session, structural images were acquired using a T1-weighted sequence (TR = 1900 ms, TE = 2.26 ms, TI = 900 ms, flip = 9°) with a resolution of 1 mm x 1 mm x 1 mm. Functional images were acquired using a gradient echo planar sequence (TR = 2000 ms, TE = 30 ms, flip angle = 80°). Each volume contained 32 axial slices which were measured in ascending order. The voxel size was 4 mm x 4 mm x 4 mm (slice gap: 20%). The experiment was controlled by E-Prime 2.0.

### 2.4 Data Analysis

#### 2.4.1 Preprocessing

We performed preprocessing using fMRIPrep 22.0.2 (O.; Esteban et al., 2019; O. Esteban et al., 2018; RRID:SCR_016216) which is based on Nipype 1.8.5 (K. J. Gorgolewski et al., 2011; K. J. Gorgolewski et al., 2018; RRID:SCR_002502).

#### 2.4.2 Anatomical data

The T1-weighted (T1w) image was corrected for intensity non-uniformity (INU) with N4BiasFieldCorrection (Tustison et al., 2010) distributed with ANTs (Avants, Epstein, Grossman, & Gee, 2008); ARRID:SCR_004757), and used as T1w-reference throughout the workflow. The T1w-reference was then skull-stripped with a Nipype implementation of the antsBrainExtraction.sh workflow (from ANTs), using OASIS30ANTs as target template. Brain tissue segmentation of cerebrospinal fluid (CSF), white-matter (WM) and gray-matter (GM) was performed on the brain-extracted T1w using fast (FSL 6.0.5.1:57b01774, RRID:SCR_002823, (Zhang, Brady, & Smith, 2001). Brain surfaces were reconstructed using recon-all (FreeSurfer 7.2.0, RRID:SCR_001847, (Dale, Fischl, & Sereno, 1999)) and the brain mask estimated previously was refined with a custom variation of the method to reconcile ANTs-derived and FreeSurfer-derived segmentations cortical gray-matter (RRID:SCR_002438, (Klein et al., 2017)).

Even though all our analyses were performed in native space, individual brains were additionally normalized to standard MNI space in order to be able to back-transform cortical network ROIs defined in MNI space back into native space (see section Cerebral Cortex Networks). To this end, volume-based spatial normalization to standard space (MNI152NLin2009cAsym) was performed through nonlinear registration with antsRegistration (ANTs 2.3.3), using brain-extracted versions of both T1w reference and the T1w template. The following template was selected for spatial normalization: *ICBM 152 Nonlinear Asymmetrical template version 2009c* ((Fonov, Evans, McKinstry, Almli, & Collins, 2009), RRID:SCR_008796; TemplateFlow ID: MNI152NLin2009cAsym).

#### 2.4.3 Functional MRI data

The MRI data from experiment Phase 2 of each subject was preprocessed as follows. First, a reference volume and its skull-stripped version were generated using a custom methodology of fMRIPrep. Head-motion parameters with respect to the reference (transformation matrices, and six corresponding rotation and translation parameters) are estimated before any spatiotemporal filtering using mcflirt (FSL 6.0.5.1:57b01774, (Jenkinson, Bannister, Brady, & Smith, 2002). MR images were slice-time corrected to 0.972s (0.5 of slice acquisition range 0s-1.95s) using 3dTshift from AFNI (Cox & Hyde, 1997); RRID:SCR_005927).

The BOLD time-series (including slice-timing correction) were resampled onto their original, native space by applying the transforms to correct for head-motion. These resampled BOLD time-series will be referred to as preprocessed BOLD in original space, or just preprocessed BOLD.

Several confounding time-series were calculated based on the preprocessed BOLD: framewise displacement (FD), DVARS and three region-wise global signals. FD was computed using two formulations following Power (absolute sum of relative motions) (Jenkinson et al., 2002; Power et al., 2014) (relative root mean square displacement between affines, Jenkinson et al. (2002)). FD and DVARS are calculated for each functional run, both using their implementations in Nipype (following the definitions by Power et al. 2014). The three global signals are extracted within the CSF, the WM, and the whole-brain masks. Additionally, a set of physiological regressors were extracted to allow for component-based noise correction (CompCor, Behzadi et al. 2007). Principal components are estimated after high-pass filtering the preprocessed BOLD time-series (using a discrete cosine filter with 128s cut-off) for the two CompCor variants: temporal (tCompCor) and anatomical (aCompCor). tCompCor components are then calculated from the top 2% variable voxels within the brain mask. For aCompCor, three probabilistic masks (CSF, WM and combined CSF+WM) are generated in anatomical space. The implementation differs from that of Behzadi et al. in that instead of eroding the masks by 2 pixels on BOLD space, a mask of pixels that likely contain a volume fraction of GM is subtracted from the aCompCor masks. This mask is obtained by dilating a GM mask extracted from the FreeSurfer’s aseg segmentation, and it ensures components are not extracted from voxels containing a minimal fraction of GM. Finally, these masks are resampled into BOLD space and binarized by thresholding at 0.99 (as in the original implementation).

Components are also calculated separately within the WM and CSF masks. For each CompCor decomposition, the k components with the largest singular values are retained, such that the retained components’ time series are sufficient to explain 50 percent of variance across the nuisance mask (CSF, WM, combined, or temporal). The remaining components are dropped from consideration. The head-motion estimates calculated in the correction step were also placed within the corresponding confounds file. The confound time series derived from head motion estimates and global signals were expanded with the inclusion of temporal derivatives and quadratic terms for each (Satterthwaite et al. 2013). Frames that exceeded a threshold of 0.5 mm FD or 1.5 standardized DVARS were annotated as motion outliers. Additional nuisance timeseries are calculated by means of principal components analysis of the signal found within a thin band (crown) of voxels around the edge of the brain, as proposed by (Patriat, Reynolds, & Birn, 2017). The BOLD reference was then co-registered to the T1w reference using bbregister (FreeSurfer) which implements boundary-based registration (Greve & Fischl, 2009). Co-registration was configured with six degrees of freedom.

The BOLD time-series were resampled onto Free Surfer’s “fsnative” space. All resamplings were performed with a single interpolation step by composing all the pertinent transformations (i.e. head-motion transform matrices and co-registrations to anatomical and output spaces). Gridded (volumetric) resamplings were performed using antsApplyTransforms (ANTs), configured with Lanczos interpolation to minimize the smoothing effects of other kernels (Lanczos, 1964). Non-gridded (surface) resamplings were performed using mri_vol2surf (FreeSurfer).

Many internal operations of fMRIPrep use Nilearn 0.9.1 (Abraham et al., 2014) (RRID:SCR_001362), mostly within the functional processing workflow (for more details of the pipeline, see the section corresponding to workflows in fMRIPrep’s documentation).

#### 2.4.4 Thalamus Segmentation

We used FreeSurfer and the thalamic probabilistic segmentation algorithm by Iglesias *et al*. (2018) which is incorporated into the FreeSurfer software. Utilising a probabilistic thalamus atlas based upon histological data, this algorithm uses Bayesian inference to segment both the left and right thalamic nuclei of individual subjects into an overall total of 47 subregions (see Figure 2 for the results of an exemplary subject). In a first step, we used SynthSeg (Billot et al., 2023) in order to obtain optimized whole-thalamus segmentation. In a second step, the thalamus was segmented into 23 subregions for the left thalamus and 24 subregions for the right thalamus (with increased stiffness of the mesh from 0.05 to 0.25), as the paracentral nucleus on the left side was not available. Here, a subset of 16 subregions per hemisphere were included in our analysis: the anteroventral (AV), central medial (CeM), centromedian (CM), lateral geniculate nucleus (LGN), lateral posterior (LP), medial geniculate nucleus (MGN), mediodorsal lateral parvocellular (MDl), mediodorsal medial magnocellular (MDm), pulvinar anterior (PuA), pulvinar medial (PuM), pulvinar lateral (PuL), pulvinar inferior (PuI), ventral anterior (VA), ventral lateral anterior (VLa), ventral lateral posterior (VLp) nuclei, and ventral posterolateral (VPL) nuclei. However, 7 subregions on the left side and 8 subregions on the right side could not be identified in all our subjects, due to their extremely small volume in the probabilistic atlas. These included the bilateral central lateral (CL), laterodorsal (LD), limitans (suprageniculate) (L-SG), medial ventral (reuniens) (MV-re), parafascicular (Pf), ventral anterior magnocellular (VAmc), ventromedial (VM), and the right paracentral (Pc) nuclei. As a result, these subregions were excluded from our analysis in all subjects.

**Figure 2.**
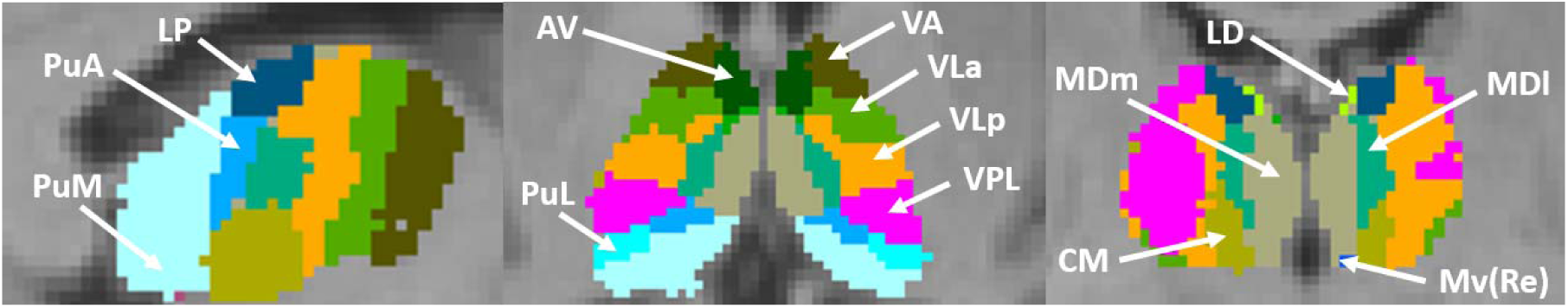
Segmentation of thalamic nuclei using an exemplary subject and a probabilistic atlas by Iglesias *et al*. (2018), displaying 13 thalamic subregions. Subregions include the following: anteroventral (AV), centromedian (CM), lateral dorsal (LD), lateral posterior (LP); mediodorsal lateral (MDl); mediodorsal medial (MDm), nucleus reuniens (Mv(Re)), pulvinar anterior (PuA), pulvinar lateral (PuL), pulvinar medial (PuM); ventral anterior (VA), ventral lateral anterior (VLa); ventral lateral posterior (VLp); and ventral posterolateral (VPL).

#### 2.4.5 Basal Ganglia and Hippocampus Segmentation

For analysis of altered thalamic connectivity with subregions of the hippocampus and components of the basal ganglia, we utilised an atlas created by (Tian, Margulies, Breakspear, & Zalesky, 2020) which segregated 38 bilateral regions (Figure 3). These ROIs were transformed from MNI space into subject-specific native space via the antsApplyTransforms function based on the MNI152NLin2009cAsym_to-T1w inverse transformation parameters created during fMRIPrep normalization.

**Figure 3.**
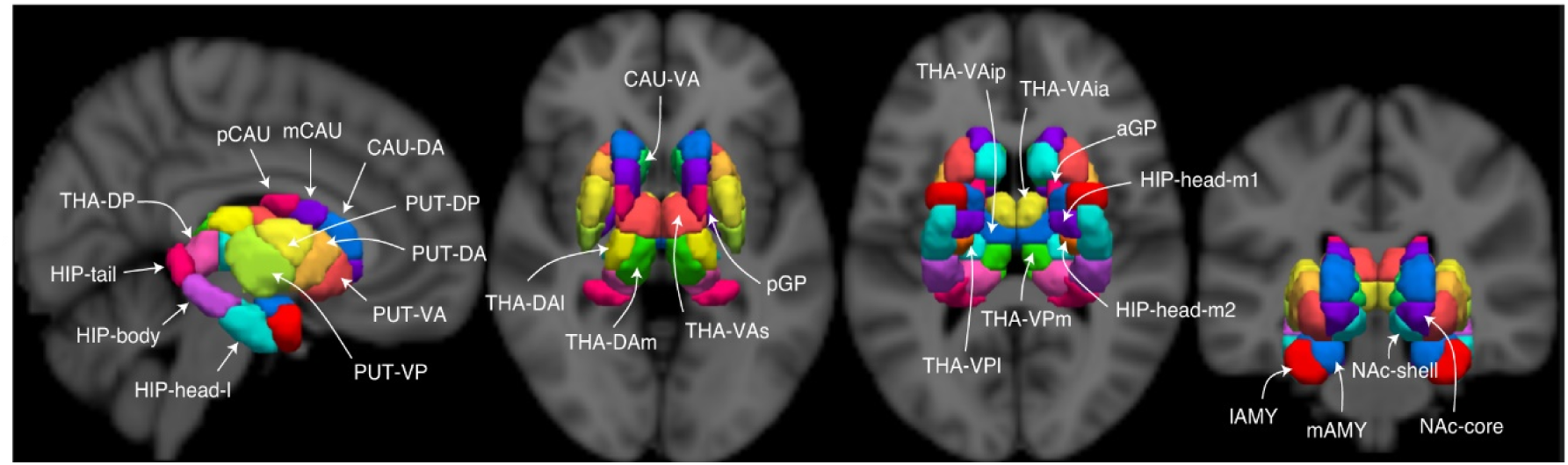
Atlas of subcortical nuclei delineating hippocampus and basal ganglia subregions. Figure adapted from Tian et al. (2020). The thalamic subregions were not utilised in our study, as thalamic segmentation was performed using the atlas from Iglesias et al. (2018) due to its provision of subject-specific segmentation and a greater number of thalamic nuclei. *Abbreviations:* Hippocampus head medial subdivisions (HIP-head-m1 and HIP-head-m2), hippocampus head lateral subdivision (HIP-head-l), hippocampus body (HIP-body), hippocampus tail (HIP-tail), putamen ventroanterior (PUT-VA), putamen dorsoanterior (PUT-DA), putamen ventroposterior (PUT-VP), putamen dorsoposterior (PUT-DP), caudate ventroanterior (CAU-VA), caudate dorsoanterior (CAU-DA), caudate medial (mCAU), caudate posterior (pCAU), lateral amygdala (lAMY), medial amygdala (mAMY), nucleus accumbens shell (NAc-shell), nucleus accumbens core (NAc-core), globus pallidus posterior (pGP), and globus pallidus anterior (GPa).

#### 2.4.6 Cerebral Cortex Networks

To analyse altered thalamic connectivity with the cerebral cortex, we utilised 12 functional network probability maps, which altogether were comprised of 153 spherical ROIs. These network ROIs were obtained within a study by Dworetsky et al. (2021) which created network templates based upon a winner-takes-all procedure from a total of six different datasets, available for download at https://github.com/GrattonLab/Dworetsky_etal_ConsensusNetworks. Here, they used an average network assignment probability of ≥ 75% to assign the 153 spherical ROIs into the available networks. These networks included those known for their roles within higher-order cognitive processes (Dworetsky et al., 2021; Hu et al., 2023; Y. Zhou et al., 2018), such as the fronto-parietal network (FPN), default mode network (DMN), salience network (SN), dorsal attention network (DAN), cingulo-opercular network (CON), parieto-medial network (PMN), parieto-occipital network (PON), and language network (which corresponds to the ventral attention network in other work by Dworetsky *et al*. (2018)) (Figure 4). To be consistent with the original paper by Dworetsky et al. (2021), we have used the label of ‘language network’ for this ROI.

**Figure 4.**
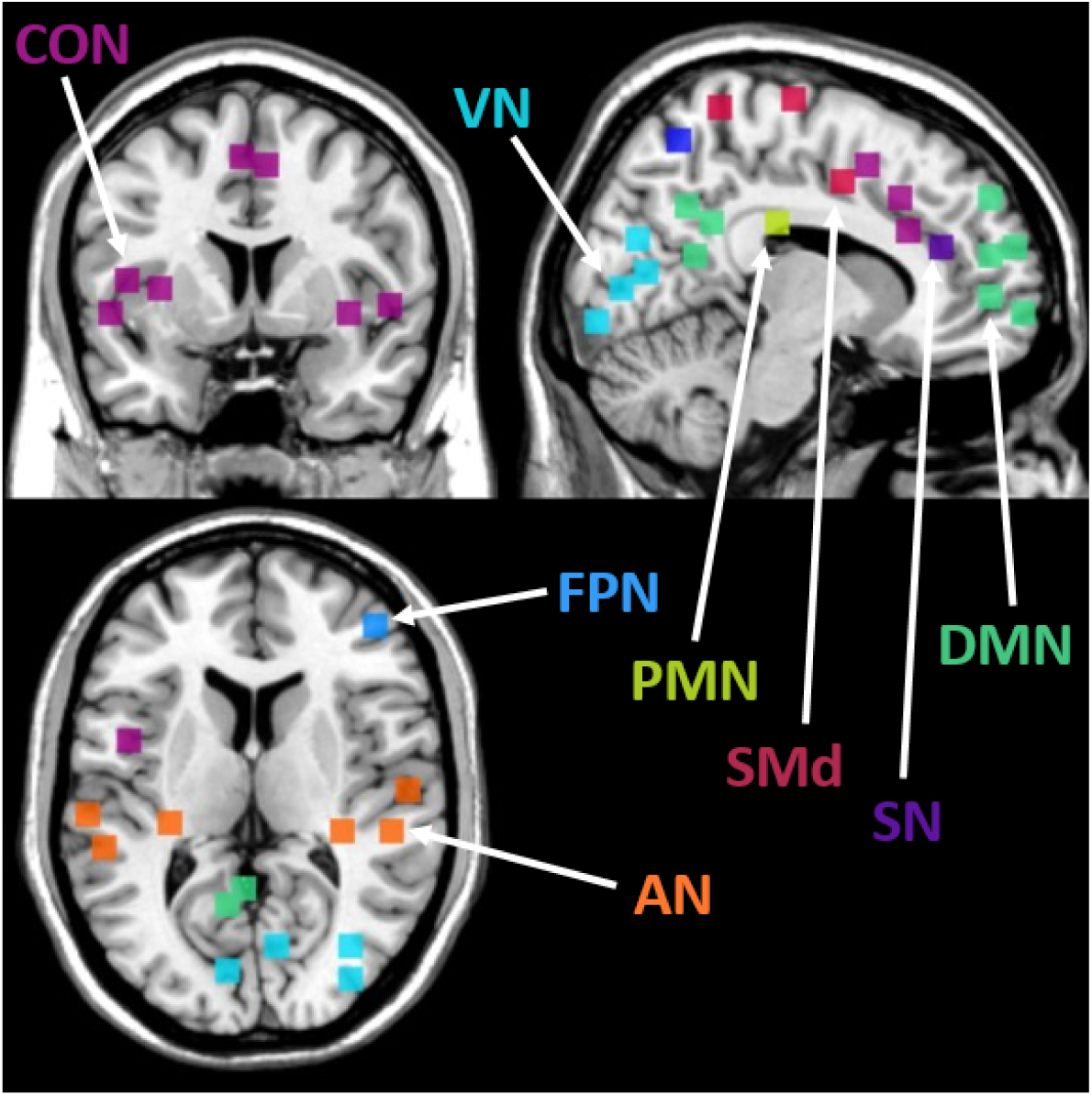
A probabilistic representation of 8 association networks from Dworetsky *et al*. (2021). Colours represent the respective networks; Auditory network, AN (orange), cingulo-opercular network, CON (purple); default mode network, DMN (green), fronto-parietal network, FPN (middle blue), parieto-medial network, PMN (yellow), somatomotor dorsal network, SMd (red), salience network, SN (dark blue), and visual network, VN (light blue). These network voxels represent brain regions with highest confidence in network assignment across subjects in the Dworetsky study.

In addition to networks more directly involved in higher order processes, visual and somatomotor dorsal (SMd) and somatomotor lateral (SMl) networks were also provided and included in our re-analysis, alongside an additional two networks, including the Temporal Pole (TPole) network and Medial Temporal Lobe (MTL) network. However, the TPole network did not reach the network assignment probability of ≥ 75% within the Dworetsky *et al*. (2018) study, and in many subjects within our own study, no voxels were found within the Medial Temporal Lobe (MTL) network. Therefore, neither of these two additional networks were included in our analysis.

The cortical network ROIs were transformed from MNI space into subject-specific native space via the antsApplyTransforms function based on the MNI152NLin2009cAsym_to-T1w inverse transformation parameters created during fMRIPrep normalization.

#### 2.4.7 Functional data analysis

##### 2.4.7.1 Denoising

After preprocessing, we first performed denoising of the functional data in native space at the single subject level. To this end, we computed a General Linear Model (GLM) for each subject using SPM12 and MATLAB 2021a. The GLM included ‘nuisance’ regressors based on a subset of the confounding variables previously extracted by fMRIPrep. The residual timeseries were saved to disk for further processing. Consistent with the fmridenoise pipleline 24HMPaCompCorSpikeReg, we included the following confounding variables. First, we included the 3 translational and 3 rotational motion parameters and their quadratic terms as well as their derivatives and the quadratic terms of the derivatives. Second, we included the first 5 principle components of the CompCor timeseries within CSF (’c_comp_cor’) and the first 5 principle components of the CompCor timeseries within white matter (’w_comp_cor’). Third, we included the DVARS timeseries and the framewise displacement (‘FD’) timeseries.

##### 2.4.7.2 Modeling task-related activity

In preparation for the functional connectivity analysis via generalized psycho-physiological interaction (gPPI) analysis, we computed for each subject a new GLM on the previously generated denoised residual timeseries to estimate task-related activity using SPM12 and additionally defining a 128 s high-pass filter. We included model regressors for correct and incorrect approach trials as well as correct and incorrect avoidance trials. In order to capture learning-related changes across correctly implemented trials, we additionally included parametric regressors that varied according to the incremental number of correctly implemented approach and avoidance trials per individual stimulus. Finally, after model estimation, we computed an ‘omnibus’ F-contrast spanning all task regressors to be used by the subsequent gPPI for extracting seed region timecourses after regressing task-related activity.

##### 2.4.7.3 Generalized PPI analysis (Whole brain parametric connectivity analysis)

Based on the GLM just described, we set up the gPPI analysis (McLaren, Ries, Xu, & Johnson, 2012) again using the GLM framework within SPM12, for each subject. To model mean task-related activity the gPPI again included all the task-related regressors defined before. Most importantly, to capture task-related functional connectivity between the seed region BOLD timecourse and the BOLD timecourses in each voxel of the brain, the gPPI model additionally included PPI regressors obtained by convolving the original task regressors with the eigenvariate timecourse determined across all voxels within the seed region. Hence, there were in total 8 additional PPI regressors for correct and incorrect approach trials as well as correct and incorrect avoidance trials plus the respective parametric change regressors. Finally, the seed region eigenvariate timecourse was added as a model regressor to model task-unrelated signal correlation between seed region and target voxel timecourses. The effect of primary interest was the parametric connectivity change across correct learning trials (approach and avoidance alike). We therefore computed for each subject and each seed region, a whole-brain contrast for the mean learning-related parametric change over approach and avoidance conditions.

##### 2.4.7.4 Functional network analysis (ROI-to-ROI connectivity)

For each subject we took the parametric gPPI contrast images for each seed ROI and extracted for each remaining ROI (the ‘target ROI’) the mean contrast value across voxels within that target ROI. Overall, this resulted in a 81×81 ROI-to-ROI connectivity matrix for each subject (i.e., 6480 pairwise connectivity changes, excluding connectivity with own seed region) which were then used as inputs for the group-level functional network analysis using the CONN toolbox (Nieto-Castanon, 2020). Note that due to the asymmetric nature of the gPPI, connectivity between ROI 1 as seed region and ROI 2 as target region was not identical with connectivity between ROI 2 as seed region and ROI 1 as target region. Hence, also the ROI-to-ROI connectivity matrices were asymmetric.

We used the CONN toolbox (Release 20.a) for the group-level statistical analysis of these ROI-to-ROI connectivity using the default Functional Network Connectivity (FNC) option. Thereby, we were able to determine which of the 6480 pairwise connectivity changes were statistically significant. FNC first organizes the ROIs into networks based upon functional connectivity similarity metrics between all ROI-to-ROI pairings using complete-linkage clustering (Sorensen, 1948). Once these networks have been defined, FNC analyses all connections between all ROI pairs, for both within- and between-network connectivity sets (Jafri, Pearlson, Stevens, & Calhoun, 2008). FNC performs a multivariate parametric General Linear Model analysis for all connections, producing an F-statistic for each pair of networks and an associated uncorrected cluster-level *p-value*. This is defined as the likelihood under the null hypothesis of a randomly-selected pair of networks showing equal or greater effects than those observed within the current pair of networks. Lastly, a FDR-corrected cluster-level p-value (Benjamini & Hochberg, 1995) is defined as the expected proportion of false discoveries amongst all network pairs with similar or larger effects across the entire set of pairs (Nieto-Castanon, 2020). Here, a cluster threshold of p < 0.05 p-*FDR* corrected was utilised, alongside a connection threshold of p < 0.05 p-uncorrected.

Lastly, we performed an analysis whereby we investigated potential correlations between the compatibility effect within Phase 3 and functional connectivity changes across all regions of interest. To this end, we used the same CONN toolbox setup as before but this time added the subject-specific compatibility effects as a covariate within the CONN toolbox multivariate GLM analysis and then tested the covariate for statistical significance regarding all connections.

#### 2.4.8 Post Hoc Analysis

Our primary analysis did not reveal significant connectivity changes involving the putamen. However, from a theoretical perspective, increasing automatization might be expected to involve the putamen. We therefore performed a post-hoc analysis with more lenient correction requirements, this time selectively focusing on altered functional connectivity between thalamic nuclei and putamen subregions (thereby reducing the number of connections to be corrected for). Moreover, we utilised the ROI-level p-FDR corrected (ROI mass/intensity) option within the CONN toolbox. In contrast to the FNC approach where correction is performed on the level of identified function networks, the post-hoc analysis applied correction on the level of ROIs. Here, a *p*-uncorrected connection threshold of 0.05 was combined with a cluster-level p-FDR corrected threshold of 0.05. Compared to the primary analysis, this approach traded increased statistical power toward putamen-related connectivity alterations against increased risk of false positives. This enabled greater chance to detect subtler connectivity changes which may have been missed with the stricter correction approach utilized for the primary analysis.

## 3 Results

### 3.1 Behavioural Results

For completeness, we include the original behavioural results as reported in Zwosta et al. (2018) for both Phase 2 and Phase 3.

In Phase 2, within-subject ANOVAs were computed for both error rates and response times across seven learning blocks and for two motivation types, namely approach versus avoid categories (Figure 5). Overall, approach and avoidance learning conditions showed similar performance improvement in error rates and response times. Error rates and response times significantly decreased across blocks as learning progressed, with *F*(6,312) = 127.84, *p* < 0.001, η2 = 0.71 and *F*(6,312) = 122.35, *p* < 0.001, η2 = 0.70, respectively. Subjects showed generally faster responses in the approach condition than in the avoidance condition (main effect motivation type *F*(1,52) = 58.93, *p* < 0.001, η2 = 0.53). A significant interaction between motivation type and learning block indicated a steeper decline in response times in the avoidance learning condition than in the approach learning condition (*F*(6,312) = 8.14, *p* < 0.001, η2 = 0.14).

**Figure 5.**
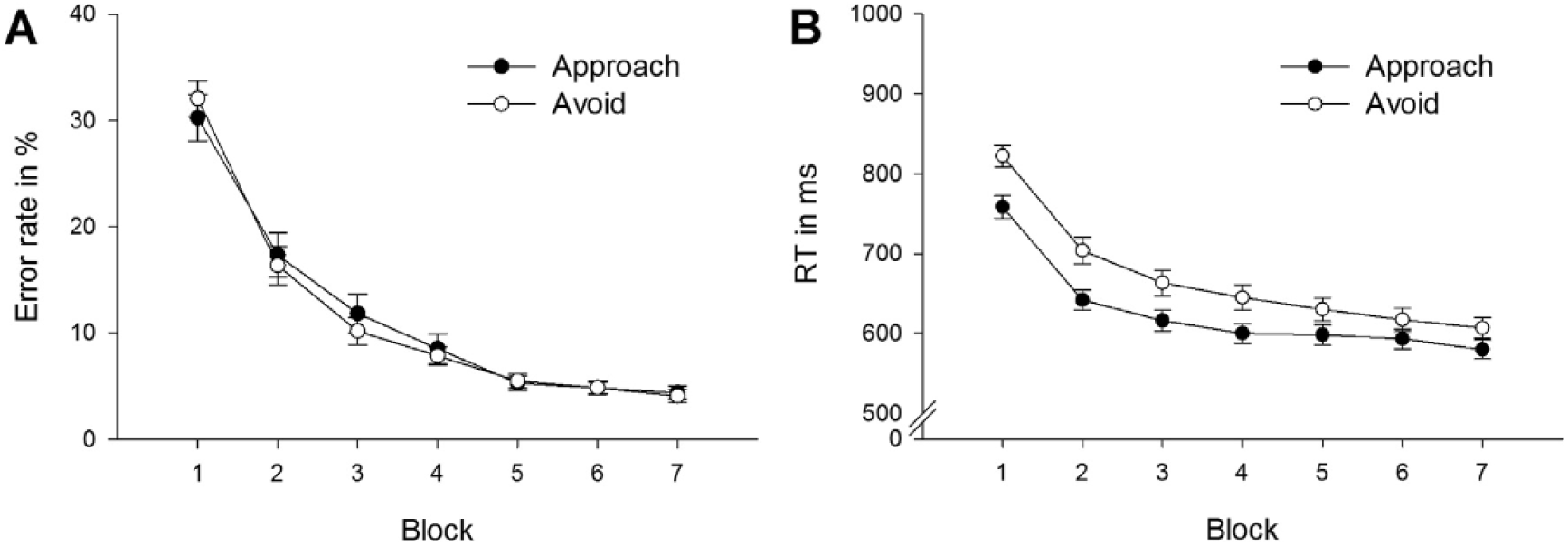
Behavioural results for Phase 2. Error rates (A) and response times (B) across learning blocks for approach and avoidance trials. Taken from (Zwosta et al., 2018). Overall, approach and avoidance learning conditions showed similar performance improvement in error rates and response times.

In Phase 3, ANOVAs were computed separately for response times and error rates, including the within-subjects factors motivation type (approach or avoid) and compatibility (compatible versus incompatible trials). Regarding error rate, there was a significant main compatibility effect in that error rates were higher for incompatible trials compared with compatible trials (*F*(1,52) = 6.58, *p* = 0.013, η^2^ = 0.11). Motivation type did not significantly influence error rates (*F*(1,52) = 0.80, p = 0.375) and there was no reliable interaction between motivation type and compatibility (*F*(1,52) = 3.20, *p* = 0.079). Regarding response times, there was also a significant main effect compatibility in that responses were slower for incompatible versus compatible trials (*F*(1,52) = 17.96, *p* < 0.001, η^2^ = 0.26). There was no main effect of motivation type (*F*(1,52) = 2.66, *p* = 0.109) nor a significant interaction effect between motivation type and compatibility (*F*(1,52) = 1.29, *p* = 0.262).

### 3.2 Learning Related Functional Connectivity Changes

Across learning trials, widespread significant alterations in functional connectivity were found between numerous thalamic nuclei and cortical networks as well as with subcortical regions (Figures 6A and 6B). For detailed statistical information regarding individual ROI-to-ROI and cluster functional connectivity alterations, please see Table 1 in the Supplementary Material.

**Figure 6.**
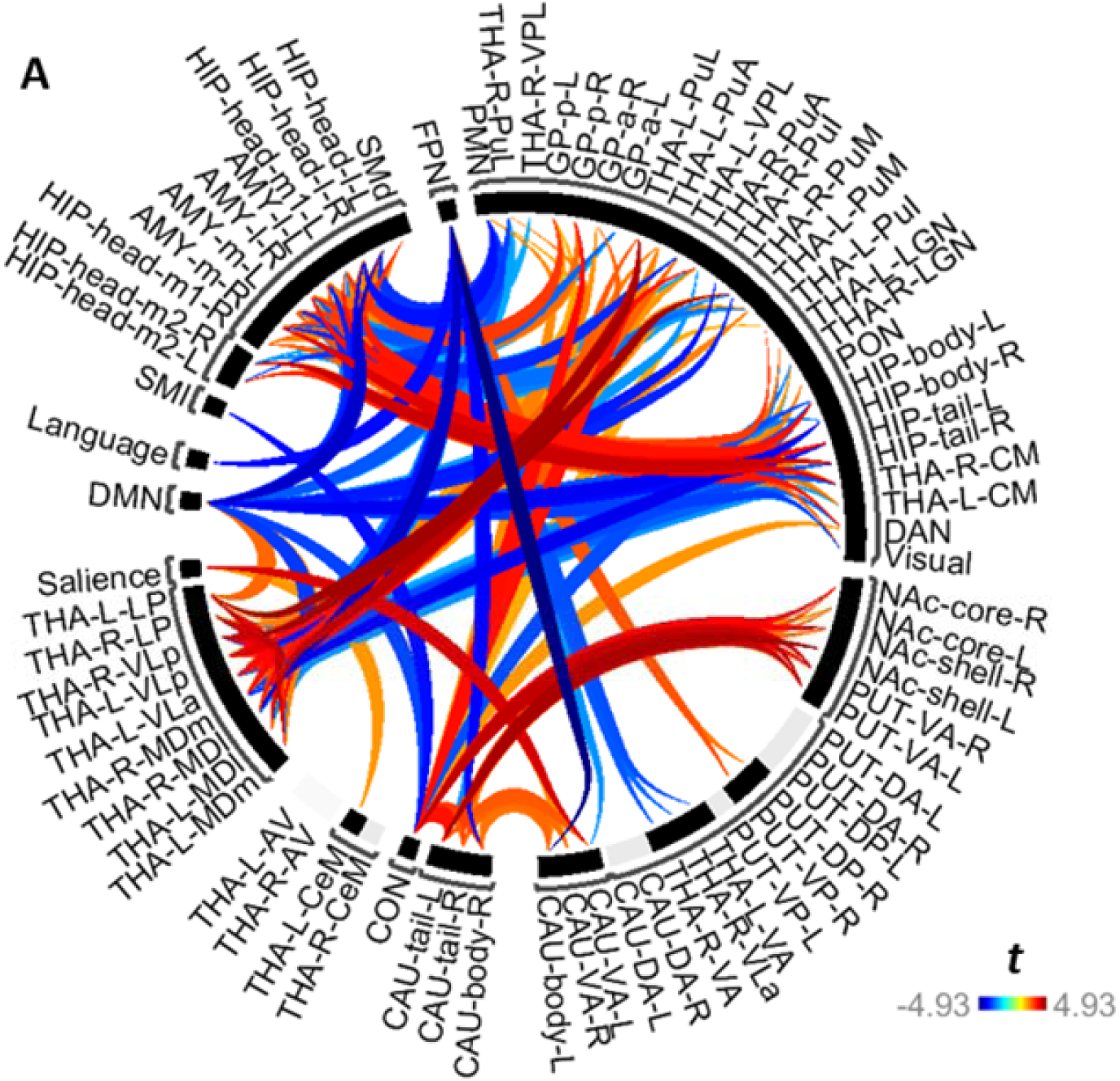
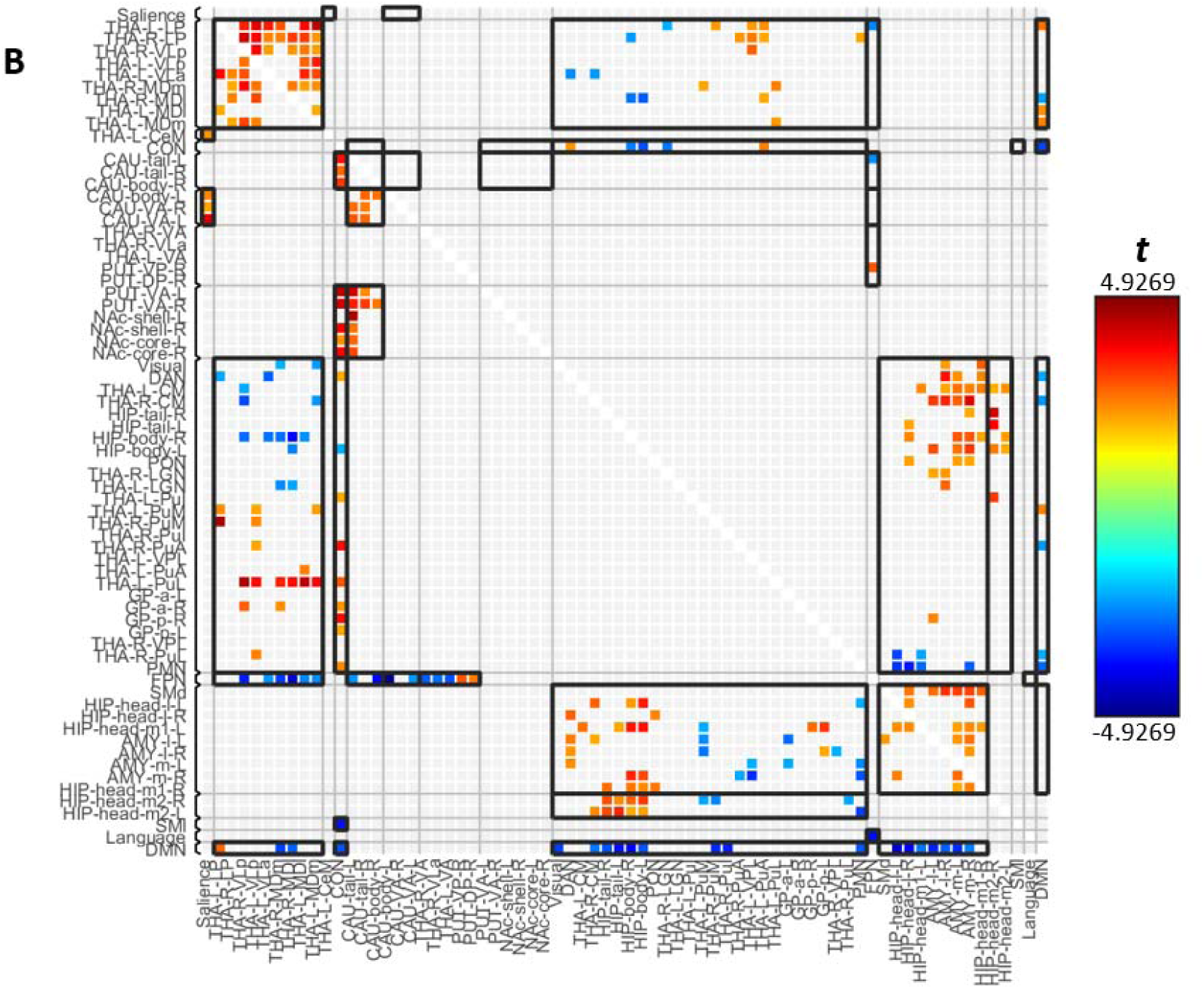
Significant functional connectivity changes amongst thalamic nuclei, non-thalamic subcortical structures, and cortical networks across learning during Phase 2 of the learning paradigm. See Supplementary Table 1 for comprehensive report of the statistical results. Blueish colours show decreasing functional connectivity, whereas reddish colours show increasing functional connectivity. Lighter colours indicate weaker effects and smaller T values (see color bar for t value range). Thin black brackets indicate cluster boundaries, as determined by the initial hierarchical clustering step during the two-stage statistical analysis within CONN. Shown are significant connectivity changes that survived multiple comparison correction based on the Functional Network Connectivity algorithm with a cluster threshold of p < 0.05 (*FDR* corrected) applied to individual connections thresholded at p < 0.05 (uncorrected). **(A)** Circular graph representation of the significant connectivity changes **(B)** Matrix display representation of the significant connectivity changes. Here, the asymmetric nature of gPPI-based connectivity values becomes visible. *Abbreviations: Organised based on 3 sets of ROIs: (1) THA, thalamus: AV, anteroventral; LGN, lateral geniculate nucleus; LP, lateral posterior; MDl, mediodorsal lateral parvocellular; MDm, mediodorsal medial magnocellular; MGN, medial geniculate nucleus; PuA, pulvinar anterior; PuI, pulvinar inferior; PuL, pulvinar lateral; PuM, pulvinar, medial; VA, ventral anterior thalamus; VLa, ventral lateral anterior; VLp, ventral lateral posterior; VPL, ventral posterolateral. (2) SUBCORTICAL: AMY, amygdala; CAU, caudate; DA, dorsoanterior; DP, dorsoposterior; GP, globus pallidus;HIP, hippocampus; NAc, nucleus accumbens; PUT, putamen; VA, ventroanterior; VP, ventroposterior. (3) CEREBRAL CORTEX: CON, cingulo-opercular; DAN, dorsal attention network; DMN, default mode network; FPN, frontoparietal network; PMN, parietal medial network; PON, parieto-occipital network; SMd, somatomotor dorsal network; and SMl, somatomotor lateral network*.

**Table 1.** List of subregions and corresponding abbreviations.

| Regions of Interest and Abbreviations |  |  |  |
| --- | --- | --- | --- |
| Thalamus | Abbreviation | Amygdala | Abbreviation |
| Anteroventral | AV | Lateral amygdala | lAMY |
| Central medial | CeM | Medial amygdala | mAMY |
| Centromedian | CM | **Caudate** |  |
| Lateral geniculate nucleus | LGN | Ventroanterior caudate | VA CAU |
| Lateral posterior | LP | Dorsoanterior caudate | DA CA |
| Mediodorsal lateral parvocellular | MDl | **Globus Pallidus** |  |
| Mediodorsal medial magnocellular | MDm | Anterior globus pallidus | aGP |
| Medial geniculate nucleus | MGN | Posterior globus pallidus | pGP |
| Pulvinar anterior | PuA | **Hippocampus** |  |
| Pulvinar inferior | PUI | Hippocampus head, medial division, subdivision 1 | HC head, medial 1 |
| Pulvinar lateral | PuL | Hippocampus head, medial division, subdivision 2 | HC head, medial 2 |
| Pulvinar medial | PuM | Hippocampus head, lateral division | HC head, lateral |
| Ventral anterior | VA | Hippocampus body | HC body |
| Ventral lateral anterior | VLa | Hippocampus tail | HC tail |
| Ventral lateral posterior | VLp | **Nucleus Accumbens** |  |
| Ventral posterolateral | VPL | Nucleus accumbens, shell | NAcc, shell |
| **Cortical Networks** |  | Nucleus accumbens, core | NAcc, core |
| Cingulo-opercular network | CON | **Putamen** |  |
| Default mode network | DMN | Ventroanterior putamen | VA PUT |
| Dorsal attention network | DAN | Dorsoanterior putamen | DA PUT |
| Frontoparietal network | FPN | Ventroposterior putamen | VP PUT |
| Language network | Language | Dorsoposterior putamen | DP PUT |
| Parietal medial network | PMN |  |  |
| Parieto-occipital network | PON |  |  |
| Salience network | Salience |  |  |
| Somatomotor dorsal network | SMd |  |  |
| Somatomotor lateral network | SMl |  |  |
| Visual network | Visual |  |  |

All results reported here are based on connectivity changes averaged across approach and avoidance learning conditions. The direct comparison between approach and avoidance learning conditions did not yield significant results. Also, we did not find significant correlations between the size of the compatibility effect and connectivity changes.

Whilst the functional connectivity changes between the ROIs in our study were clustered based upon similarity metrics within the present data set, this also coincided with known functional properties regarding cortical and subcortical regions in previous literature as described below (Figures 6A-B).

This means the ROIs within our study tended to form clusters with other subregions of the same region, such as thalamic nuclei clustering with many other thalamic subregions (especially those of a known similar functional role), hippocampal subregions having a tendency to cluster together (Strange, 2022), as well as amygdala subregions (Keshavarzi, Sullivan, Ianno, & Sah, 2014). Furthermore, basal ganglia subregions also tended to form clusters amongst their own subregions, including the globus pallidus, caudate and putamen (Mattfeld, Gluck, & Stark, 2011). Furthermore, most cortical networks were clustered independently from each other. Most notably, higher order thalamic nuclei formed a cluster which consisted of many known CSTC loop nodes, namely MD, VLa, and VLp which together comprise anterior cingulate, dorsolateral frontal, lateral orbitofrontal, and motor CSTC loops (Sugiyama et al., 2018).

For convenience, we also display the connectivity alterations for selective ROI clusters (defined by the initial hierarchical clustering step) in separate figures below (Figures 7A-7D). Overall, the results were predominantly driven by decreasing functional connectivity involving three cortical networks (frontoparietal, cingulo-opercular, and default mode networks) with almost exclusively higher order thalamic nuclei, including mediodorsal, ventrolateral, lateral posterior, pulvinar, and ventral anterior subdivisions (Figures 7A-C). In addition, learning was also associated with increasing *intrathalamic* functional connectivity amongst a cluster of higher order nuclei and decreasing intrathalamic nuclei between these higher order nuclei with LGN and CM regions (Figure 7D).

**Figure 7.**
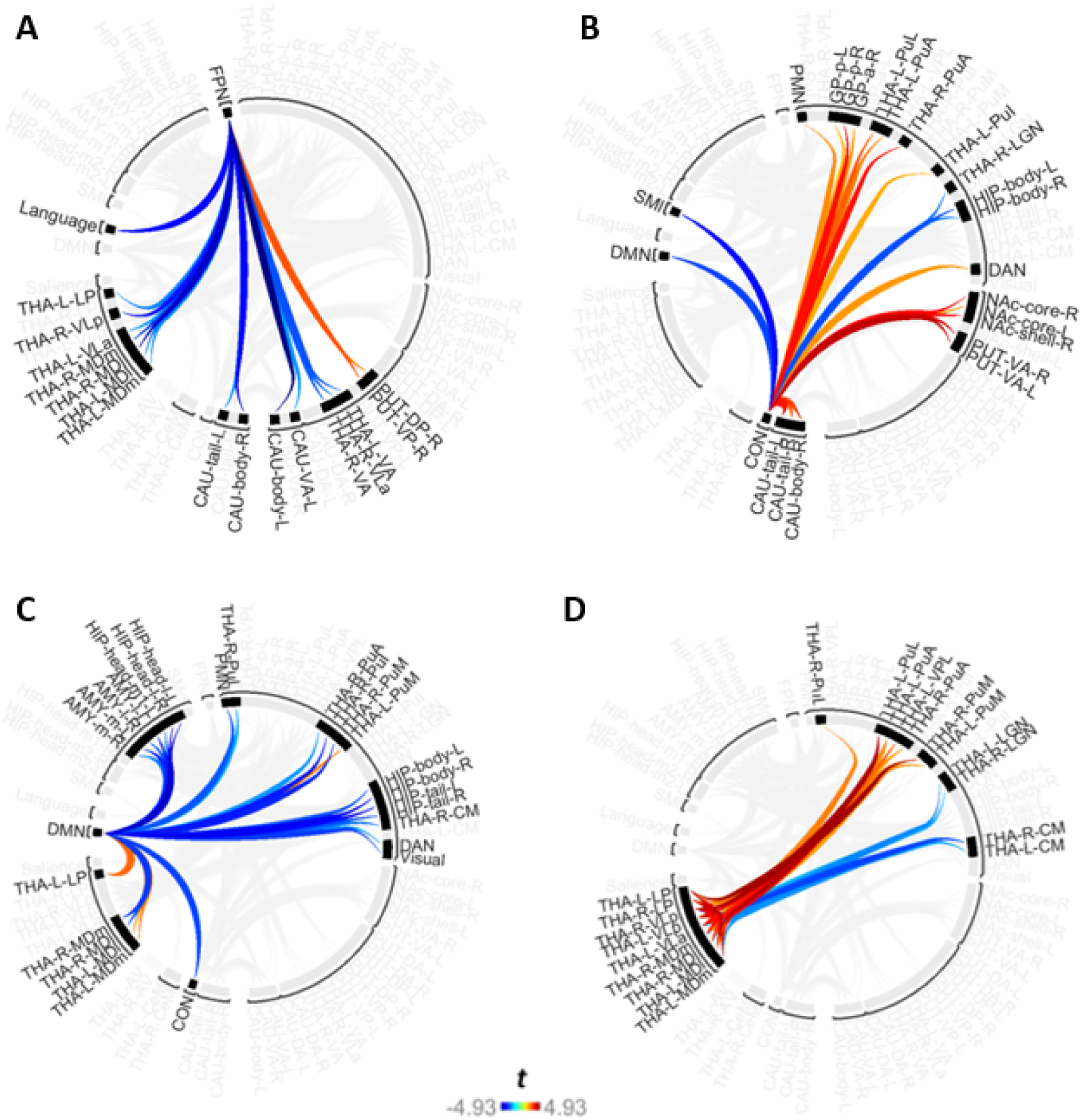
Functional connectivity alterations between thalamic regions and cortical networks and amongst thalamic nuclei across learning. This is a selective visual representation of the full results shown in Fig. 6. The selected clusters include (A) functional connectivity alterations between the frontoparietal network and thalamic regions, (B) functional connectivity changes between the cingulo-opercular network and thalamic regions, (C) functional connectivity changes between the DMN network and mediodorsal thalamic regions, and (D) functional connectivity alterations amongst thalamic regions. Red brackets indicate cluster boundaries which have been determined by the hierarchical clustering algorithm.

#### 3.2.1 Changes in connectivity during learning in a frontoparietal-thalamic network (Figure 7A)

The FPN exhibited widespread decreasing functional connectivity alterations with thalamic nuclei across learning trials (Figure 7A). Notably, all of these thalamic nuclei were higher order nuclei, most of which serve as proposed or established nodes within CSTC networks (Aton, 2021; Provost, Hanganu, & Monchi, 2015), namely the mediodorsal, ventral anterior, ventral lateral, and lateral posterior nuclei.

Together, these results suggest that across learning, functional connectivity decreases between FPN and higher order thalamic nuclei as the influence of cognitive control processes over behaviour diminishes whilst behaviour becomes more automatic.

#### 3.2.2. Alterations in connectivity during learning in a thalamo-cingulo-opercular network (Figure 7B)

In contrast, the CON network primarily showed increasing functional connectivity with various pulvinar nuclei across learning trials (Figure 7B). These alterations occurred between CON and bilateral PuA, left PuI, and left PuL regions. Conversely, the only decrease in functional connectivity that was found occurred between CON and the right LGN. Notably, all of these thalamic nuclei are regions of the visual thalamus (Purushothaman, Marion, Li, & Casagrande, 2012), with the pulvinar being primarily a higher order nucleus, whilst the LGN is a first order nucleus (Sherman, 2007).

Overall, this functional reorganisation of CON with visual thalamic regions across successful repetitions may therefore shift control away from the FPN and towards CON across learning. This would suggest that cortical networks contribute differentially to goal-directed behaviour as control transitions toward automaticity.

#### 3.2.3 Connectivity changes during learning in a thalamo-default-mode network (Figure 7C)

The DMN showed widespread altered functional connectivity with numerous thalamic regions (Figure 7C), almost all of which were either higher order nuclei or those involved in CSTC networks (Aton, 2021; Provost et al., 2015; Saalmann, 2014; Sherman, 2007). These results were totally lateralised, whereby right MD, right pulvinar, and right CM nuclei showed decreasing functional connectivity with the DMN, whilst the left MD, left PuM, and left LP showed increasing functional connectivity with the DMN.

These results thereby suggest that the thalamus shows lateralised functional differences regarding its contributions to the DMN across learning transition from goal-directed to habitual behaviour, and specifically that both MD and pulvinar nuclei are particularly important for the DMN across this transition.

#### 3.2.4 Altered intrathalamic connectivity during learning (Figure 7D)

Finally, widespread changes in functional connectivity was observed amongst numerous thalamic nuclei (Figure 7D). Most notably, these alterations featured increasing functional connectivity amongst the same higher order nuclei and CSTC nodes which exhibited altered connectivity with FPN, CON, and DMN networks. Specifically, these included MD, pulvinar, VLa, VLp, and LP nuclei. Conversely, decreases in functional connectivity were observed between these higher order thalamic nuclei and the bilateral CM and first order LGN.

These results therefore suggest that as automaticity increases through learning, higher order thalamic nuclei functionally reorganise by decreasing their functional connectivity with the FPN, increasing their functional connectivity with higher order visual thalamic regions, and increase their functional connectivity amongst themselves, potentially forming an intrathalamic network in facilitation of automatic behaviour. This may also be supported by the lateralised effects within the thalamus, whereby left MD and pulvinar regions show increasing functional connectivity with the DMN, whilst those on the right side show decreasing functional connectivity.

#### 3.2.5 Connectivity changes between thalamic nuclei and non-thalamic subcortical ROIs (hippocampus and amygdala) (Figures 8A-B)

In addition to functional connectivity alterations between cortical networks and thalamic ROIs, numerous functional connectivity alterations occurred between thalamic nuclei and non-thalamic subcortical ROIs. With the exception of two ROI-to-ROI connectivity changes between thalamic nuclei and the anterior globus pallidus, all functional connectivity alterations occurred between thalamic nuclei with hippocampus and amygdala regions.

First, functional connectivity alterations between thalamic nuclei and hippocampal subregions revealed widespread decreasing connectivity across all higher order and CSTC node thalamic nuclei, namely CM, LP, MD, pulvinar, and VL subregions, spanning 19 ROI-to-ROI connections (Figure 8A). Conversely, only one increase in functional connectivity was observed, and occurred between left PuI and right hippocampal head, medial division 2.

**Figures 8A-B.**
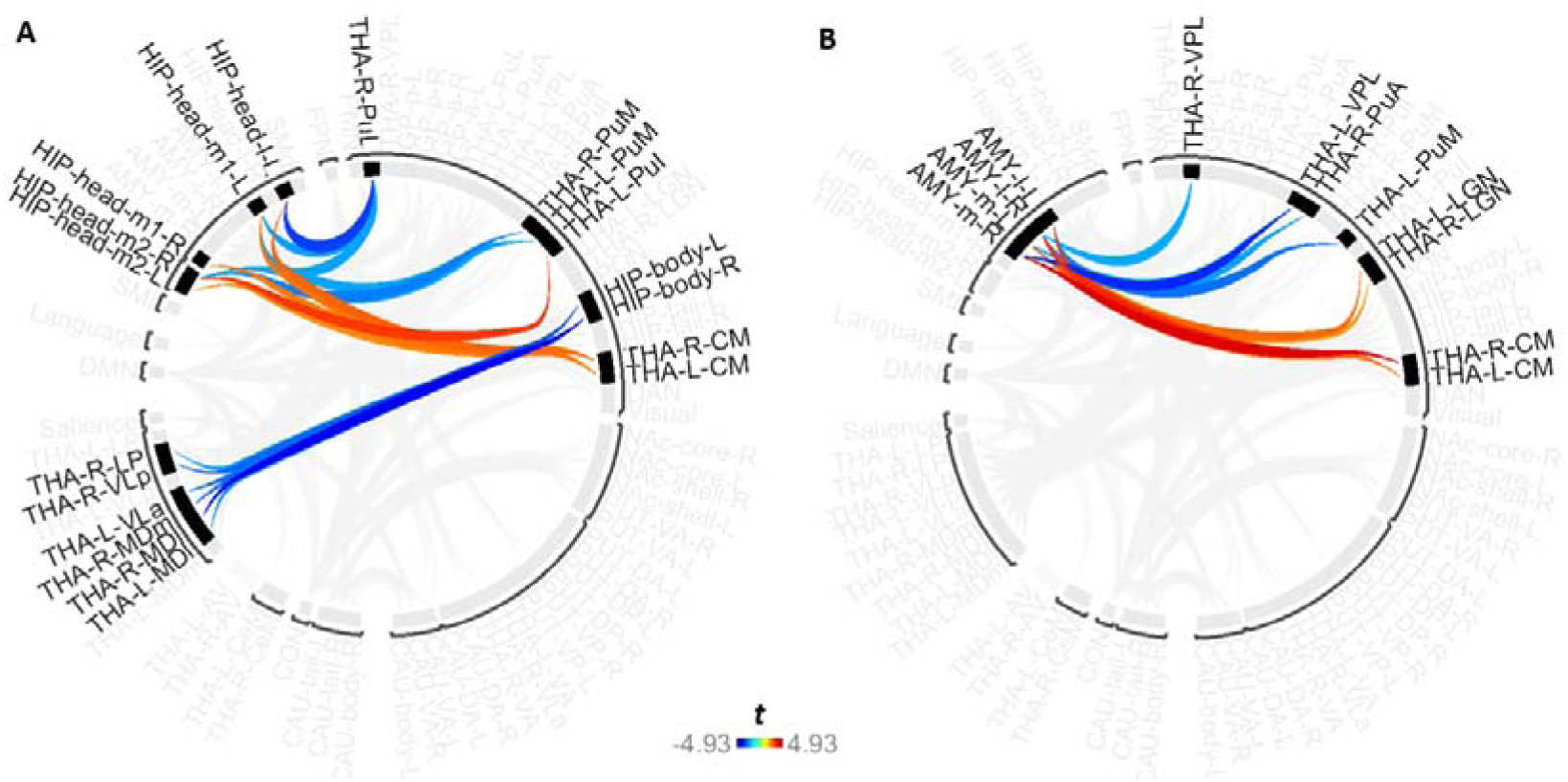
Functional connectivity alterations between thalamic nuclei with hippocampal and amygdala subregions across learning. This is a selective visual representation of the full results shown in Fig. 6. Shown are functional connectivity alterations between thalamic and hippocampal subregions (8A), and functional connectivity alterations between thalamic and amygdala subregions (8B).

Second, functional connectivity alterations also occurred between thalamic nuclei and amygdala subregions. Consistent with findings regarding the hippocampus, widespread decreasing connectivity was also observed between thalamic nuclei with amygdala subregions. However, this did not primarily occur amongst higher order or CSTC node nuclei, as this instead occurred amongst CM, LGN, and VPL, alongside two pulvinar subregions (Kim, Shin, Wang, & Nowakowski, 2023; Saalmann, 2014). This suggests that the amygdala may play a separate functional contribution to this learning transition across goal-directed toward automatic behaviour, as mediated by its altered connectivity with lower order sensory and consciousness regions of the thalamus. Furthermore, the various amygdala subregions also showed increasing functional connectivity with each other as behaviour became increasingly automatic.

Altogether, this suggests that not only does the thalamus decrease or reorganise its functional connectivity with cortical networks, but that the thalamus also diminishes functional connectivity with other subcortical areas across the transition from goal-directed to automatic behaviour within learning. Collectively, this appears to decrease communication with cortical networks, alongside generating increasing communication amongst increasingly segregated subcortical regions, such as amongst the thalamus, hippocampus, and amygdala, as learning progresses.

### 3.3 Post-Hoc Analysis

The Post Hoc analysis revealed significantly altered functional connectivity between numerous thalamic nuclei and putamen subregions, see Figure 9 and Supplementary Table 2. Here, various thalamic nuclei showed functional connectivity changes with putamen subregions, however these were restricted to two subregions, namely ventroanterior and dorsoposterior regions. The ventroanterior putamen showed increasing functional connectivity with bilateral MDm, MDl, and VLp subregions, alongside increasing functional connectivity with left AV and right LP nuclei. Conversely, the dorsoposterior putamen exhibited both increasing functional connectivity with bilateral AV and decreasing functional connectivity with right CM and PuA nuclei.

**Figures 9A-C.**
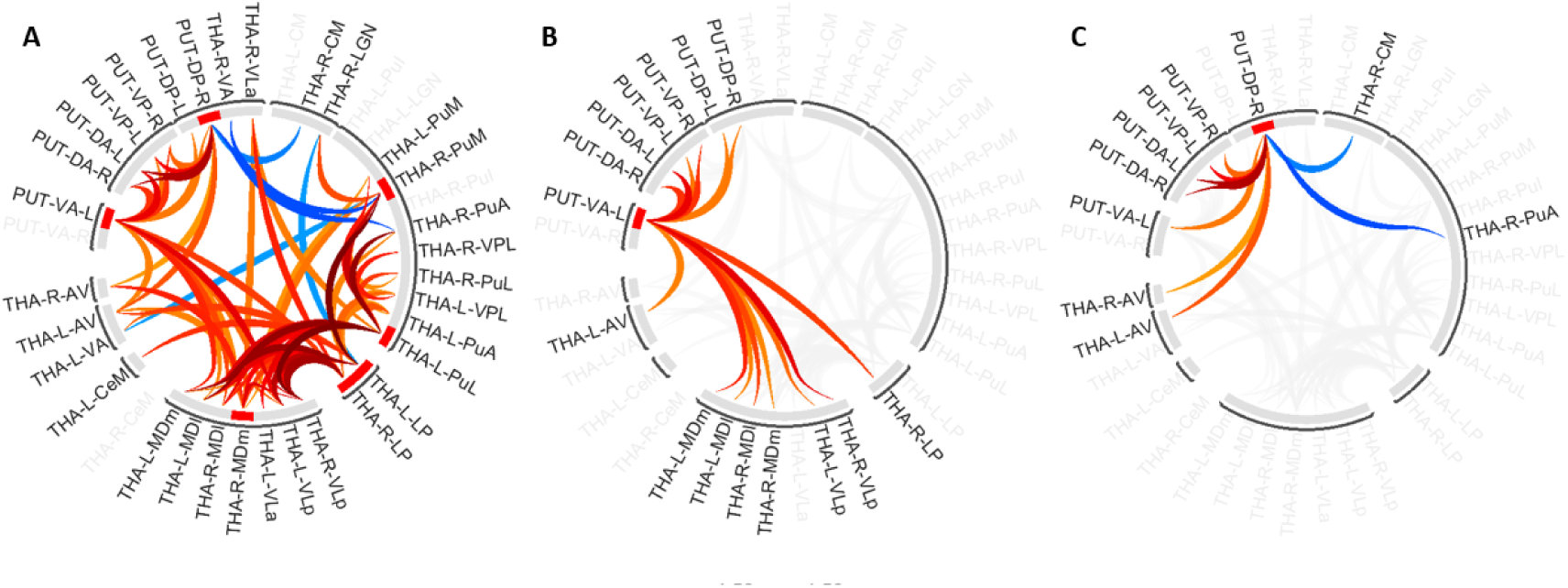
Significant functional connectivity alterations between thalamic nuclei and putamen subregions across learning. See Supplementary Table 2 for comprehensive report of the statistical results. All significant functional connectivity alterations between thalamic nuclei and putamen subregions (9A), alongside selective visual representations of the full results within Figure 9A, for the ventroanterior putamen and thalamic nuclei (9B), and for the dorsoposterior putamen with thalamic nuclei (9C). Note that these results were obtained by a post-hoc analysis selectively focusing on connections between thalamus subregions and putamen subregions with more lenient statistical correction requirements as the primary analysis.

Overall, this suggests that during learning, increasing automaticity is associated with altered functional connectivity between thalamic nuclei with putamen subregions, specifically putamen subregions which are involved in sensorimotor processing, alongside the encoding of stimulus value and stimulus-driven motivation (Graff-Radford, Williams, Jones, & Benarroch, 2017).

## 4 Discussion

Our results showed that during the transition from goal-directed toward automatic behaviour, numerous thalamic nuclei exhibited functional connectivity alterations with cerebral cortex networks, other thalamic nuclei, and subcortical regions. These changes primarily included decreasing functional connectivity between FPN and higher order thalamic regions, increasing functional connectivity between CON and the higher order visual thalamus, and lateralised functional connectivity changes between higher order thalamic nuclei and the DMN, alongside increased intrathalamic functional connectivity. Lastly, increasing functional connectivity also occurred amongst subcortical subregions, with increased functional segregation of these same subcortical subregions as behaviour became increasingly automatic.

### 4.1 Functional Reorganisation Occurs Between Numerous Cortical Networks and Higher Order Thalamic Nuclei as Learning Progresses

Goal-directed behaviour is coordinated by a variety of cortical networks, which include executive networks such as CON and FPN (X. Wang et al., 2024; Wood & Nee, 2023). These networks have diverse contributions to behaviour, suggesting not only that these networks should show altering levels of activation across the transition from goal-directed toward habitual behaviour, but also that these networks should show altering functional connectivity with thalamic nuclei which also have specialised contributions to goal-directed behaviour, namely higher order thalamic nuclei or CSTC nodes.

Accordingly, our results indeed show that the FPN, CON, and DMN exhibit learning-related changes in functional connectivity with various thalamic nuclei, and that these primarily occurred amongst higher order thalamic nuclei. These changes were also consistent with previously reported findings regarding the roles of higher order thalamic nuclei within goal-directed behaviour, as outlined in further detail in the following subsections.

### 4.2 Task-Positive Networks Show Differential Functional Connectivity Patterns Across Learning

The successful completion of a cognitive task requires the activation of various cortical networks, which are therefore known as task-positive networks (Mills et al., 2018), and include FPN, CON and DAN (Yu et al., 2019). In accordance with this, our results revealed that a variety of task-positive networks showed altered functional connectivity with thalamic nuclei across learning. However, our results showed that these alterations primarily occurred between higher order thalamic nuclei with FPN and CON networks. Notably, the thalamus is a known node of both the FPN (Camilleri et al., 2018; Dixon et al., 2018) and CON (Sadaghiani & D’Esposito, 2015).

First, the FPN showed widespread decreases in functional connectivity with higher order nuclei or CSTC nodes, whilst the CON showed increasing functional connectivity with solely the higher order visual thalamus (namely pulvinar nuclei), whilst decreasing functional connectivity with the first order visual thalamus (the LGN). Here, the FPN serves as a crucial task-positive network alongside the CON (Sadaghiani & D’Esposito, 2015), whereby the function of the FPN is to facilitate flexible and goal-directed behaviour using feedback from responses (Hausman et al., 2022). On the other hand, the role of the CON is to implement task sets (Marek & Dosenbach, 2018) and to regulate actions (D’Andrea et al., 2023). In this way, it has been suggested that the FPN initiates and adjusts cognitive control, whilst the CON maintains the task set (Zanto & Gazzaley, 2013).

Consequently, the altered functional connectivity between the FPN and CON with higher order thalamic nuclei within our results appear to align with the known functional roles of these regions across goal-directed toward habitual behaviour. It may therefore be the case that goal-directed behaviour is initially driven by strong connections between the FPN and higher order thalamic nuclei, enabling responses to be outcome-driven, and that this high connectivity becomes progressively diminished as automaticity forms. Additionally, the increasing functional connectivity between the CON and pulvinar during our visual task may have aided successful implementation of the task set of visual S-R associations, as this increase in functional connectivity between CON and pulvinar nuclei also aligns with the pulvinar’s specialised role in handling visual input (Berman & Wurtz, 2008). Here, implementation of the task set is sustained through the transition from goal-directed to habitual behaviour (Schneider & Logan, 2014), a function which is present even within habitual behaviour and in its early phases may reflect “strategic automaticity” (Sheeran, Webb, & Gollwitzer, 2006).

Together, this lends further support to the differential contributions of cortical networks within the coordination of goal-directed versus habitual behaviour, and suggests that the generation of behaviour across this behavioural transition is mediated by functional reorganisation with select higher order thalamic nuclei. Consequently, it may be the case that executive cortical networks may have a hierarchical organisation within goal-directed behaviour, whereby transition to habit is characterised by reduced contributions of executive networks which first begin with the FPN, whilst the CON sustains its influence upon behaviour across a longer period and toward automaticity. This would be consistent with findings that higher order cognitive control has a gradient from higher-to-lower or rostral-to-caudal regions (Choi, Drayna, & Badre, 2018), such that within the frontal lobe, a hierarchy flows from schematic control toward contextual and finally sensorimotor control (Zhu & Han, 2022). Accordingly, the nodes of the FPN and CON fall within this hierarchy, as does the rostral-caudal organisation of higher order versus first order nuclei within the thalamus (Vertes, Linley, Groenewegen, & Witter, 2015).

### 4.3 The DMN shows Diverging Functional Connectivity Changes with Lateralised Thalamic Subregions across Learning

The generation of automatic behaviour is associated with DMN activation (X. Zhou & Lei, 2018) and de-coupling of the DMN with task-positive cognitive networks (Baumann, Schäfer, & Ruge, 2023; Vatansever, Menon, & Stamatakis, 2017). Here, in contrast to task-positive cognitive networks, task-negative networks activate during internally-directed attention, and include the DMN (S. Weber, Aleman, & Hugdahl, 2022). The DMN is involved in internally-focused thought processes, such as mind-wandering, future planning, self-reflection and memory recall (Menon, 2023), and it is known to activate during distraction and boredom, especially within repetitive tasks (Danckert & Merrifield, 2018). Repetitive tasks, by nature, foster habitual responding (Stojanovic, Fries, & Grund, 2021). Consistent with this, our own results also show that during increasing automaticity, decoupling occurred between the DMN and task-positive cortical networks, including CON, DAN, and PMN.

Furthermore, this functional decoupling occurred alongside altered functional connectivity with lateralised thalamic regions. Here, left-hemispheric thalamic regions showed increasing functional connectivity with the DMN, whilst right-hemispheric thalamic regions showed decreasing functional connectivity. These changes specifically included occurred amongst higher order nuclei, namely the left MDl, MDm, LP and PuL, and the right MDl, MDm, PuA, PuI, and PuM. This is aligned with a huge body of research which reports lateralization across the cerebral cortex regarding many cognitive and executive functions, including decision making (Corser and Jasper, 2014), attention (Bartolomeo and Seidel Malkinson, 2019), and in the access of semantic information (Liuzzi, Ubaldi, & Fairhall, 2021). Therefore, our results suggest that higher order thalamic nuclei also have lateralised functional contributions to goal-directed and habitual behaviour, and is consistent with a growing body of research which reports thalamic functional lateralisation across sensorimotor functions (Herrero, Barcia, & Navarro, 2002), language (Llano, 2013), and speech (D. Wang et al., 2022). There is also evidence to suggest that the DMN shows some lateralisation in its functioning, and that even with age, the DMN is mostly lateralised to the left side and gradually shifts toward rightward lateralisation (Banks et al., 2018). However, the functional significance of this lateralisation of the DMN as a network, and in terms of its functional connectivity with the MD thalamus, is unclear.

### 4.4 The Mediodorsal Thalamus is Involved in Switching from the FPN Toward the DMN across learning

The MD thalamus has been identified as a special thalamic region within goal-directed behaviour. Functionally, it appears to be central in coordinating flexible responses (J. Weber et al., 2023) through the updating of action-outcome and stimulus-outcome associations across rule or contingency changes (M. Wolff & Vann, 2019), as well as in the maintenance of rule representations (Schmitt et al., 2017). This ability is essential within the flexible control of behaviour and within our own learning paradigm, in order for subjects to successfully discard associations learnt within previous phases and to maintain relevant and current task rules for successful performance across task phases and within each phase. Therefore, the MD nuclei appear to have critical roles in response-outcome driven behaviour.

Here, the MD nuclei are known to connect with the FPN and support its functioning within cognitive tasks (Li et al., 2022; Shine, Lewis, Garrett, & Hwang, 2023), and they are also argued to be a central node of the DMN, thereby also influencing DMN functioning during internally focused cognition (Aguilar & McNally, 2022). Furthermore, the MD nuclei are classically defined by their extensive connections with the prefrontal cortex (PFC) (Delevich, Tucciarone, Huang, Li, & Id, 2015), for which the PFC is a region which is also well known for being central in generating flexible goal-directed behaviour (J. Weber et al., 2023). Therefore, as what might be expected, the MD nucleus accordingly projects primarily to the prefrontal cortex and is considered to be the ‘principal thalamic nucleus for’ (Zikopoulos & Barbas, 2007) and ‘essential partner of [the] prefrontal cortex’ (Parnaudeau, Bolkan, & Kellendonk, 2018). Here, the MD nuclei are known nodes within both the prefrontal or executive CSTC loop (Sugiyama et al., 2018) as well as the limbic CSTC loop (Gibson et al., 2017), for which they also connect extensively with the basal ganglia (J. B. Smith, Smith, Venance, & Watson, 2022). At the level of the thalamus, the MD nuclei serve as the ‘associative nucleus of the thalamus’ (Ji et al., 2019), consistent with the prefrontal lobe as an association area which mediates top-down processing of sensory and motor information (Siddiqui, Chatterjee, Kumar, Siddiqui, & Goyal, 2008; Spellman, Svei, Kaminsky, Manzano-Nieves, & Liston, 2021). Finally, extensive connections also run to and from other association cortices and the limbic system, enabling the MD nuclei to serve as an integration centre for somatosensory information (Hika & Khalili, 2023). Here, cortical networks, including the FPN and DMN, form from these association areas (Yeo et al., 2011) in order to drive behaviour.

Therefore, the MD nuclei are in a unique and prime position to coordinate cortical network activity, including amongst task-positive cognitive networks such as the FPN and the task-negative DMN, and may thereby serve as a mechanism for switching behavioural control across task-positive versus task-networks. Here, through this dynamic interaction between MD nuclei with both the FPN versus DMN, the reorienting of attention between external versus internal loci of attention (V. Smith, Mitchell, & Duncan, 2018) could take place across shorter timescales or within early learning, facilitating flexible behavioural switching between response-outcome versus stimulus-outcome associations (Parnaudeau et al., 2018). Through this mechanism, the MD nuclei may therefore mediate the attention switching which occurs between the external task and internal phenomena during behavioural switching throughout learning, and ultimately, on a longer time frame, this could also underlie the transition from goal-directed toward automatic control of behaviour, via reduced functional connectivity with the FPN and stabilised functional connectivity with the DMN, thereby driving goal-directed versus automatized control of behaviour, respectively.

As such, we propose that the MD nuclei may therefore serve as a critical mechanism for switching between goal-directed versus automatized behaviour. Here, we speculate that the MD thalamus may act as a crucial thalamic node across cortical networks, through its increasing functional connectivity with the FPN during cognitive tasks and in its switching of connectivity instead toward the DMN during automaticity. This would thereby enable the MD nuclei to contribute to cognitive tasks and then to monitor automatic behaviours as behaviour becomes increasingly automatized.

### 4.5 Subcortical and Cortical Systems Become Increasingly Functionally Segregated as Automaticity Forms

A growing body of research is reporting that habitual behaviour is driven predominantly by subcortical regions of the brain, with cortical areas diminishing their influence as goal-directed contributions decrease (Gillan, Robbins, Sahakian, van den Heuvel, & van Wingen, 2016; Liljeholm, Dunne, & O’Doherty, 2015). In accordance with this, our own results show decreasing functional connectivity between several cortical networks and widespread regions of the thalamus and other subcortical subregions, which included FPN, DAN, PMN, the hippocampus, and amygdala. Together, this suggests that as behaviour becomes increasingly automatic, that cortical and subcortical regions become increasingly functionally segregated and isolated. Here, response-outcome contingencies become progressively devalued or removed altogether as habits come to fruition (B. Gardner, 2015; Schreiner, Renteria, & Gremel, 2020), with behaviour becoming increasingly driven by stimuli instead of outcome. Overall, these functional connectivity changes appear to reflect decreasing cortical influence upon behaviour, and a shift towards subcortical control with the thalamus at its centre; a region which continues to receive bottom-up information from the sensory periphery (Mashour & Hudetz, 2017). Consequently, this may give rise to the impaired awareness and reflection upon behaviour which arises with automaticity (Benjamin Gardner et al., 2024).

Furthermore, these same subcortical regions also became increasingly functionally segregated from each other, with the exception of the nucleus accumbens and caudate nucleus. This was also the case between the thalamus and hippocampus, and suggests that as behaviour becomes more automatic, the thalamus also becomes increasingly functionally isolated from memory processing areas, alongside cognitive areas. However, conversely, functional connectivity increased amongst these same subcortical nuclei, such that subregions of the caudate nucleus, hippocampus and amygdala became increasingly connected with subregions of their own anatomical region.

Altogether, this aligns with the vast body of research which suggests that the subcortical system becomes increasingly permitted to drive responses without moderation by cortical networks, driving transition from outcome-driven towards stimulus-driven responses, even when doing so produces errors as a result of rule or contextual changes (K. S. Smith & Graybiel, 2016). Here, this delegation of behavioural responding to the subcortical system in automaticity has been hailed as a ‘fundamental innovation in human cognition’ (Shine & Shine, 2014) which frees the cerebral cortex to handle more complex and abstract psychological mentation, through this delegation of control to lower order areas. At the connectivity level, this behavioural change may therefore be mediated through a transition from task-positive networks via elevated thalamocortical connectivity toward increased control amongst increasingly functionally segregated subcortical structures and decreased thalamocortical connectivity, giving rise to habitual control.

### 4.6 A Potential Intrathalamic Network is Associated with Increasing Automaticity

As aforementioned, not only does transition occur from cortical toward subcortical control, but this is mirrored by transition toward a less connected thalamocortical system, as the thalamus diminishes connectivity with FPN and DAN, and reorganises its activity with DMN. However, in addition to increasing functional segregation across learning and automaticity, higher order thalamic nuclei and CSTC nodes showed increasing functional connectivity amongst each other, namely between LP, MDl, MDm, PuA, PuL, PuM, VLa, and VLp nuclei. Consequently, it may be the case that as automaticity develops, thalamic nuclei thereby potentially form a network specific for mediating or assisting the generation of automatic behaviour. Here, the exception of increasing functional connectivity between the CON and pulvinar is consistent, due to the CON’s role in implementing task sets (Dosenbach et al., 2006), and potentially within strategic automaticity before automaticity eventually solidifies and becomes controlled via thalamic rather than cortical processes.

Here, the thalamus is therefore a prime candidate to assume control over behaviour in automaticity. It is a highly unique subcortical region which is formed from the posterior part of the forebrain within the embryo (Scholpp & Lumsden, 2010), and has been termed “the principal information hub” of the vertebrate brain (Govek et al., 2022). Here, the thalamus receives bottom-up input from almost all of the senses (Courtiol & Wilson, 2014), integrates multisensory information (Tyll, Budinger, & Noesselt, 2011), generates consciousness (Whyte, Redinbaugh, Shine, & Saalmann, 2024), and generates motor responses in coordination with other subcortical areas (Bosch-Bouju, Hyland, & Parr-Brownlie, 2013), as cortical motor areas reduce their influence within habitual movements or after long term learning (E. J. Hwang et al., 2019). Altogether, this thereby potentially reflects an evolutionarily older neural system prior to the development of the cerebral cortex, with the thalamus at the apex.

In support of this, research reports a network of intrathalamic pathways across the thalamus. In humans, the anterior and MD nuclei together comprise the limbic thalamus and together are nodes within the memory system (M. Wolff, Alcaraz, Marchand, & Coutureau, 2015), the anterior nuclei connect to the laterodorsal (LD) nuclei, an interconnected complex exists between the LP and pulvinar nuclei, whilst the LGN in turn connects to this complex (Casanova & Chalupa, 2023; Rodrigo-Angulo & Reinoso-Suárez, 1988). In addition, complexes are known to form between VA and VL nuclei within CSTC loops, and also between centromedian and parafascicular nuclei (Johnson et al., 2020). Connectivity is also reported amongst dorsal thalamic nuclei as well as amongst intralaminar nuclei in rodents and cats, respectively (Crabtree & Isaac, 2002), the LD nuclei connect with the LGN in monkeys, cats, and rodents (Perry & Mitchell, 2019), and it is estimated that the rodent thalamus has over 2,000 intrathalamic connections in each side of the thalamus, with thalamic nuclei also showing connectivity with its homologous nucleus on the other brain hemisphere (Swanson, Sporns, & Hahn, 2019). However, arguably the most prominent example of intrathalamic connectivity is that of the thalamic reticular nucleus in humans and animals (Viviano & Schneider, 2015), which wraps around other thalamic nuclei and provides inhibitory control over them in order to selectively modulate their thalamocortical connections (Halassa et al., 2014).

Furthermore, subdivisions of individual nuclei are also intuitively likely to be connected with each other given their contributions to similar functional domains. For example, pulvinar subdivisions (anterior, inferior, lateral, and medial) which are involved in visual or multi-sensory processes (Kaas & Lyon, 2007), as well as anterior (anteroventral, anterodorsal, and anteromedial) and MD (lateral and medial) subdivisions which contribute to memory and cognitive processes, or both, respectively (de Kloet et al., 2021; Mathiasen, O’Mara, & Aggleton, 2020).

In support of this, recent findings have been reported in schizophrenia that altered intrathalamic functional connectivity is observed between dorsolateral, ventral anterior, and ventromedial portions of the thalamus (Gong et al., 2019). This is of particular interest given the impaired goal-directed behaviour that can be associated with this disorder (Bowie & Harvey, 2006; Cooper et al., 2019), including in their impaired action-outcome learning, inflexible responses and reduced behavioural adaptation (Morris, Cyrzon, Green, Le Pelley, & Balleine, 2018). Altogether, this disorder is therefore indicative of crucial intrathalamic functional connectivity patterns which likely contribute to normal functioning within healthy subjects, and may contribute to the dysfunction that is observed in schizophrenia. Here, further research could utilise the learning paradigm within our study, alongside a sample of patients with schizophrenia, to investigate functional connectivity abnormalities within this cohort. In this, it could be expected that alterations in functional connectivity patterns across the transition from goal-directed behaviour toward automaticity will be found, given the impairments in task shifting in schizophrenia alongside its status as a disorder of connectivity amongst large-scale networks, which specifically reflect reduced network integration and segregation (Y. Wang, Hu, & Li, 2022). Indeed, whilst such investigation is sparse, at least one study has investigated this and found abnormal circuitry underlying habitual behaviour in schizophrenic subjects (Weickert et al., 2002), with others also proposing that these abnormalities in the habit system exist (Avery et al., 2019; Morris et al., 2018). Further research in light of this could provide valuable insight into this matter.

### 4.7 Thalamic Nuclei Show Altered Functional Connectivity with Putamen Subregions

From a theoretical perspective, increasing automatization might be expected to involve the putamen. However, our primary analysis did not reveal significant connectivity changes involving the putamen. In light of the theoretical relevance, we performed a post-hoc analysis with more lenient correction requirements. This investigation of altered functional connectivity selectively between thalamic nuclei and putamen subregions unveiled changes occurring between primarily higher order thalamic nuclei with ventroanterior and dorsoposterior subregions of the putamen. Here, the ventroanterior putamen exhibited increasing functional connectivity with bilateral MDm, MDl, and VLp, alongside left AV and right LP nuclei. Moreover, the dorsoposterior putamen showed increasing functional connectivity with bilateral AV and decreasing functional connectivity with right PuA and CM nuclei.

The putamen is a component of the basal ganglia system, serving alongside the caudate nucleus as the dorsal striatum and main input region to the basal ganglia (Zheng et al., 2023). The putamen receives excitatory projections from cortical sensorimotor, midline thalamus and cerebellar areas (Kunimatsu, Maeda, & Hikosaka, 2019; Simioni, Dagher, & Fellows, 2016; Starr et al., 2011), and serves as a node within both direct and indirect pathways for motor generation and inhibition, respectively (Young, Reddy, & Sonne, 2023). Therefore, the putamen acts as an action filter for behaviour within goal-directed behaviour (Hikosaka et al., 2019) and is also well-known for its importance in generating habitual behaviour (Balleine & O’Doherty, 2010; Tricomi, Balleine, & O’Doherty, 2009).

Results from a meta-analysis reveal a range of functional differences between anterior and posterior subdivisions of the putamen (Pauli, O’Reilly, Yarkoni, & Wager, 2016). While both sub-regions are related to sensorimotor processes, the anterior putamen is functionally connected not only with sensorimotor regions in the cerebral cortex, but also appears to have language-related specialisations. In contrast, the posterior putamen is connected not only to cortical sensorimotor areas, but also with temporal cortex and insula regions. Furthermore, recent research finds that the anterior putamen is critical for newly acquired habits, whereas more extensively trained stimulus-response associations seem to rely more heavily on the dorsoposterior putamen (Guida, Michiels, Redgrave, Luque, & Obeso, 2022).

Therefore, within our own study, transition from goal-directed toward habitual control may have been facilitated by these thalamus-putamen functional connectivity changes. In accordance with this, these nuclei included higher order thalamic nuclei, alongside thalamic nuclei which are established nodes of direct and indirect basal ganglia pathways (Mitchell, 2015; Perry & Mitchell, 2019; Rocha et al., 2023). This suggests that thalamic nuclei may mediate the transition from goal-directed to more habitual behaviour through their connections not only with goal-directed cortical networks, but also through their dynamic connectivity with putamen subregions.

### 4.8 No Significant Association Between Functional Connectivity Changes and Behaviour

Previous results in humans showed a rather mixed picture regarding the association between neural and behavioural markers of habit learning (Gera et al., 2023; Tricomi et al., 2009; X. Wang, Zwosta, Wolfensteller, & Ruge, 2023; Zwosta et al., 2018). It has been notoriously difficult to reliably find such an association with respect to the putamen as one of the prime candidate brain regions typically assumed to be involved in automatic control based on animal research (Gera et al., 2023; Zwosta et al., 2018). The present study adds to this picture, as there was no significant correlation between connectivity changes and the size of the compatibility effect as a putative proxy of acquired habit strength. Overall, caution seems advised regarding the validity of behavioural markers of habit strength, at least in the context of human behaviour (Pool et al., 2022; Watson & de Wit, 2018).

### 4.9 Results for larger nuclei, but higher resolution needed for smaller nuclei

In the present study, we utilised a relatively low spatial resolution of 4 mm x 4 mm x 4 mm. This resolution is adequate to distinguish distinct thalamic nuclei across subjects such as the relatively large MD and pulvinar (Byne et al., 2002; Highley, Walker, Crow, Esiri, & Harrison, 2003). For some smaller nuclei, such as lateral dorsal or parafascicular nuclei (Mai & Majtanik, 2018), or subsections within MD, this resolution may not allow for accurately distinguishing thalamic nuclei. Within our own study, the mean nuclei sizes for individual MD nuclei subregions ranged from 253 mm³ to 817 mm³. Furthermore, numerous other thalamic nuclei also ranged in size within these functional voxel limits, see Supplementary Table 1. However, for clearer delineation of thalamic nuclei, and the inclusion of smaller thalamic nuclei, higher resolution MRI studies are therefore necessary. In addition, our mean LGN size was 250 mm, which is larger than typically reported, and may be due to a poor T1 image signal affecting lateral portions of the thalamus.

## 5 Conclusions

Across learning and as automaticity increases, thalamic nuclei exhibit various functional connectivity alterations with cerebral cortex networks and non-thalamic subcortical regions. These alterations occurred predominantly amongst higher order thalamic nuclei, for which there are three key conclusions. First, we found that thalamic nuclei exhibit decreasing functional connectivity with cerebral cortex networks, especially the FPN as automaticity increases. Second, we observed that the DMN exhibit decoupling with other cerebral cortex networks, for which the mechanism for switching between FPN and DMN may be mediated by MD thalamic nuclei. Third, automaticity was characterised by increasing functional connectivity amongst higher order thalamic nuclei, which may therefore constitute an intrathalamic network. Altogether, this indicates that learning is characterised by massive functional reorganisation of the thalamocortical system, resulting in functional segregation of cerebral cortex versus subcortical systems as behaviour becomes increasingly supported by subcortical structures. Finally, this may underlie the decreased influence of cognition and increasingly stimulus-driven mode of behaviour which characterises automatic action.

