## Supplementary Material for "Progressive Changes in Functional Connectivity between Thalamic Nuclei and Cortical Networks Across Learning"

Supplementary Materials

**Supplementary Table 1.** Detailed results for the group-level functional connectivity analysis obtained using the Functional Network Connectivity multivariate parametric statistics implemented in the CONN toolbox. The table lists significant connectivity clusters for network pairs with p <0.05 *FDR*-corrected across clusters (in bold font) together with univariate statistics for individual connections within each cluster surviving an uncorrected threshold of p < 0.05.

| **Analysis Unit** | **Statistic** | **p-unc** | **p-FDR** |
| --- | --- | --- | --- |
| **Cluster 1/130** | **F(2,50) = 12.83** | 0.000032 | 0.004144 |
| Connection PUT-VA-L – CON | T(51) = -4.28 | 0.000082 | 0.005083 |
| Connection PUT-VA-R – CON | T(51) = -4.16 | 0.000122 | 0.009516 |
| Connection NAc-R – CON | T(51) = -3.74 | 0.000463 | 0.036119 |
| Connection NAc-shell-R – CON | T(51) = -3.72 | 0.000495 | 0.038585 |
| Connection NAc-core-L – CON | T(51) = -2.26 | 0.027800 | 0.317209 |
| **Cluster 2/130** | **F(2,50) = 11.34** | 0.000087 | 0.005658 |
| Connection FPN – CAU-body – L | T(51) = -4.93 | 0.000009 | 0.000719 |
| Connection FPN – CAU-VA – L | T(51) = -2.27 | 0.027682 | 0.093877 |
| **Cluster 3/130** | **F(2,50) = 9.56** | **0.000305** | **0.010507** |
| Connection NAc-shell-L – CAU-tail-L | T(51) = 4.51 | 0.000039 | 0.003018 |
| Connection PUT-VA-L – CAU-tail-L | T(51) = 4.14 | 0.000130 | 0.005083 |
| Connection PUT-VA-R – CAU-tail-L | T(51) = 3.44 | 0.001171 | 0.045660 |
| Connection PUT-VA-R – CAU-tail-R | T(51) = 3.12 | 0.002963 | 0.057771 |
| Connection NAc-core-R – CAU-tail-L | T(51) = 2.99 | 0.004329 | 0.104600 |
| Connection PUT-VA-L – CAU-tail-R | T(51) = 2.36 | 0.022241 | 0.108550 |
| Connection PUT-VA-R – CAU-body-R | T(51) = 2.55 | 0.013978 | 0.136289 |
| Connection NAc-shell-R – CAU-tail-L | T(51) = 2.62 | 0.011541 | 0.225059 |
| Connection NAc-core-L – CAU-tail-L | T(51) = 2.73 | 0.008744 | 0.227345 |
| **Cluster 4/130** | **F(2,50) = 9.48** | **0.000323** | **0.010507** |
| Connection Thal-L-LP – Thal-L-MDm | T(51) = 4.32 | 0.000072 | 0.005590 |
| Connection Thal-L-LP – Thal-L-VLp | T(51) = 4.02 | 0.000195 | 0.007594 |
| Connection Thal-R-LP – Thal-R-VLp | T(51) = 4.21 | 0.000103 | 0.008016 |
| Connection Thal-R-LP – Thal-L-VLp | T(51) = 3.79 | 0.000401 | 0.015649 |
| Connection Thal-L-LP – Thal-L-MDl | T(51) = 3.62 | 0.000675 | 0.017538 |
| Connection Thal-L-LP – Thal-L-VLa | T(51) = 3.44 | 0.001163 | 0.022029 |
| Connection Thal-L-LP – Thal-R-VLp | T(51) = 3.38 | 0.001412 | 0.022029 |
| Connection Thal-R-VLp – Thal-L-VLp | T(51) = 3.76 | 0.000438 | 0.034128 |
| Connection Thal-R-MDm – Thal-R-VLp | T(51) = 3.67 | 0.000577 | 0.045009 |
| Connection Thal-R-LP – Thal-L-MDl | T(51) = 3.15 | 0.002755 | 0.046021 |
| Connection Thal-R-LP – Thal-R-MDl | T(51) = 3.12 | 0.002950 | 0.046021 |
| Connection Thal-L-VLa – Thal-L-LP | T(51) = 3.66 | 0.000594 | 0.046314 |
| Connection Thal-L-LP – Thal-R-MDm | T(51) = 2.88 | 0.005775 | 0.048789 |
| Connection Thal-L-VLa – Thal-L-MDl | T(51) = 3.34 | 0.001580 | 0.061633 |
| Connection Thal-L-MDm – Thal-R-VLp | T(51) = 2.99 | 0.004245 | 0.078893 |
| Connection Thal-L-VLp – Thal-L-MDm | T(51) = 3.41 | 0.001272 | 0.099188 |
| Connection Thal-L-VLa – Thal-L-MDm | T(51) = 3.02 | 0.003888 | 0.101097 |
| Connection Thal-L-VLa – Thal-R-VLp | T(51) = 2.90 | 0.005543 | 0.108093 |
| Connection Thal-R-LP – Thal-R-MDm | T(51) = 2.43 | 0.018498 | 0.144288 |
| Connection Thal-R-LP – Thal-L-VLa | T(51) = 2.38 | 0.021070 | 0.148926 |
| Connection Thal-R-LP – Thal-L-MDm | T(51) = 2.26 | 0.027829 | 0.148926 |
| Connection Thal-L-VLa – Thal-R-LP | T(51) = 2.41 | 0.019664 | 0.153379 |
| Connection Thal-L-VLp – Thal-L-MDl | T(51) = 2.77 | 0.007745 | 0.159490 |
| Connection Thal-L-VLp – Thal-R-VLp | T(51) = 2.60 | 0.012268 | 0.159490 |
| Connection Thal-L-MDm – Thal-L-VLp | T(51) = 2.44 | 0.018416 | 0.159604 |
| Connection Thal-R-MDl – Thal-L-VLp | T(51) = 3.01 | 0.004079 | 0.196501 |
| Connection Thal-R-VLp – Thal-L-MDl | T(51) = 2.51 | 0.015327 | 0.199250 |
| Connection Thal-R-MDl – Thal-R-LP | T(51) = 2.43 | 0.018743 | 0.219273 |
| Connection Thal-R-MDm – Thal-L-VLp | T(51) = 2.57 | 0.013128 | 0.222907 |
| Connection Thal-R-MDm – Thal-R-MDl | T(51) = 2.52 | 0.014924 | 0.222907 |
| Connection Thal-R-MDm – Thal-L-MDm | T(51) = 2.32 | 0.024662 | 0.240452 |
| Connection Thal-R-VLp – Thal-R-MDl | T(51) = 2.24 | 0.029425 | 0.286895 |
| Connection Thal-R-VLp – Thal-L-VLa | T(51) = 2.18 | 0.033649 | 0.288506 |
| Connection Thal-R-VLp – Thal-L-MDm | T(51) = 2.14 | 0.036988 | 0.288506 |
| Connection Thal-L-MDm – Thal-R-LP | T(51) = 2.02 | 0.048526 | 0.290494 |
| Connection Thal-R-MDm – Thal-L-MDl | T(51) = 2.12 | 0.039050 | 0.292776 |
| Connection Thal-R-MDm – Thal-R-LP | T(51) = 2.03 | 0.047420 | 0.308228 |
| Connection Thal-L-MDl – Thal-L-MDm | T(51) = 2.05 | 0.045421 | 0.557397 |
| Connection Thal-L-MDl – Thal-L-LP | T(51) = 2.03 | 0.047075 | 0.557397 |
| **Cluster 5/130** | F(1,51) = 13.23 | 0.000642 | 0.011985 |
| Connection Language – FPN | T(51) = -3.64 | 0.000642 | 0.025037 |
| **Cluster 6/130** | F(2,50) = 8.45 | 0.000690 | 0.011985 |
| Connection SMd – AMY-l-R | T(51) = 3.23 | 0.002171 | 0.077454 |
| Connection SMd – AMY-m–L | T(51) = 3.14 | 0.002780 | 0.077454 |
| Connection SMd – AMY-m-R | T(51) = 3.08 | 0.003309 | 0.077454 |
| Connection SMd – HIP-head-m1-R | T(51) = 2.76 | 0.008053 | 0.104691 |
| Connection SMd – AMY-l-L | T(51) = 2.67 | 0.010078 | 0.112297 |
| Connection AMY-l-L – AMY-m-R | T(51) = 2.62 | 0.011491 | 0.112692 |
| Connection SMd – HIP-head-l-R | T(51) = 2.53 | 0.014443 | 0.118568 |
| Connection AMY-m-R – AMY-m-L | T(51) = 2.68 | 0.009886 | 0.128520 |
| Connection HIP-head-m1-L – AMY-m-R | T(51) = 2.35 | 0.022917 | 0.130747 |
| Connection HIP-head-m1-L – HIP-head-R | T(51) = 2.34 | 0.023467 | 0.130747 |
| Connection AMY-m-R – HIP-head-l-L | T(51) = 2.46 | 0.017424 | 0.135906 |
| Connection HIP-head-m1-L – HIP-head-l-L | T(51) = 2.28 | 0.026970 | 0.140243 |
| Connection AMY-l-L – AMY-m-L | T(51) = 2.24 | 0.029818 | 0.147354 |
| Connection HIP-head-m1-L – HIP-head-m1-R | T(51) = 2.18 | 0.034073 | 0.147400 |
| Connection HIP-head-m1-L – AMY-m-L | T(51) = 2.16 | 0.035905 | 0.147400 |
| Connection AMY-l-L – SMd | T(51) = 2.20 | 0.032272 | 0.148072 |
| Connection HIP-head-m1-R – AMY-m-R | T(51) = 2.46 | 0.017214 | 0.200160 |
| Connection HIP-head-l-L – AMY-m-R | T(51) = 2.87 | 0.005976 | 0.222108 |
| Connection HIP-head-m1-R – AMY-m-L | T(51) = 2.03 | 0.048036 | 0.288213 |
| Connection HIP-head-l-L – HIP-head-l-R | T(51) = 2.27 | 0.027186 | 0.358960 |
| Connection AMY-l-R – AMY-m-R | T(51) = 2.25 | 0.028628 | 0.366128 |
| **Cluster 7/130** | F(2,50) = 8.36 | 0.000736 | 0.011985 |
| Connection Thal-L-PuL – Thal-R-VLp | T(51) = 4.53 | 0.000036 | 0.001759 |
| Connection Thal-L-PuL – Thal-L-MDl | T(51) = 4.34 | 0.000068 | 0.001759 |
| Connection Thal-R-PuM – Thal-L-LP | T(51) = 4.56 | 0.000032 | 0.002507 |
| Connection Thal-L-PuL – Thal-L-MDm | T(51) = 3.71 | 0.000507 | 0.007914 |
| Connection Thal-L-PuL – Thal-L-VLp | T(51) = 3.60 | 0.000720 | 0.008410 |
| Connection Thal-L-PuL – Thal-R-MDl | T(51) = 3.58 | 0.000755 | 0.008410 |
| Connection Thal-L-PuL – Thal-R-MDm | T(51) = 3.40 | 0.001299 | 0.012662 |
| Connection HIP-body-R – Thal-R-MDl | T(51) = -3.84 | 0.000341 | 0.026600 |
| Connection Thal-R-CM – Thal-R-VLp | T(51) = -3.03 | 0.003884 | 0.060584 |
| Connection Thal-R-LP – Thal-L-VPL | T(51) = 2.51 | 0.015183 | 0.133682 |
| Connection Thal-L-LP – Thal-L-PuA | T(51) = 2.30 | 0.025820 | 0.143852 |
| Connection HIP-body-R – Thal-R-VLp | T(51) = -2.72 | 0.008975 | 0.144805 |
| Connection HIP-body-R – Thal-L-VLa | T(51) = -2.67 | 0.010270 | 0.144805 |
| Connection HIP-body-R – Thal-R-MDm | T(51) = -2.63 | 0.011139 | 0.144805 |
| Connection Thal-R-LP – Thal-R-PuA | T(51) = 2.24 | 0.029566 | 0.148926 |
| Connection Thal-R-LP – HIP-body-R | T(51) = -2.22 | 0.030549 | 0.148926 |
| Connection Thal-L-VLa – DAN | T(51) = -2.41 | 0.019521 | 0.153379 |
| Connection HIP-body-L – Thal-R-MDl | T(51) = -2.51 | 0.015448 | 0.156478 |
| Connection Thal-L-LP – Thal-R-PuM | T(51) = 2.20 | 0.032659 | 0.164003 |
| Connection Thal-L-LP – Thal-L-VPL | T(51) = 2.18 | 0.033642 | 0.164003 |
| Connection HIP-body-R – Thal-L-MDl | T(51) = -2.35 | 0.022706 | 0.177110 |
| Connection Thal-R-LP – PMN | T(51) = 2.11 | 0.039523 | 0.181341 |
| Connection DAN – Thal-L-VLa | T(51) = -2.79 | 0.007403 | 0.192474 |
| Connection Thal-R-MDl – HIP-body-L | T(51) = -2.87 | 0.005984 | 0.196501 |
| Connection Thal-R-MDl – HIP-body-R | T(51) = -2.67 | 0.010077 | 0.196501 |
| Connection Thal-R-LP – Thal-L-PuA | T(51) = 2.05 | 0.045649 | 0.197814 |
| Connection Thal-R-VLp – Thal-L-VPL | T(51) = 2.76 | 0.008072 | 0.199250 |
| Connection Thal-L-VLa – Thal-R-CM | T(51) = -2.24 | 0.029761 | 0.202949 |
| Connection Thal-L-LP – Thal-R-LGN | T(51) = -2.03 | 0.048114 | 0.205976 |
| Connection aGP-R – Thal-R-VLp | T(51) = 2.75 | 0.008243 | 0.207863 |
| Connection aGP-R – Thal-R-MDm | T(51) = 2.28 | 0.027108 | 0.207863 |
| Connection Thal-R-MDm – Thal-L-PuL | T(51) = 2.46 | 0.017147 | 0.222907 |
| Connection Thal-L-MDm – Thal-L-PuL | T(51) = 2.17 | 0.034817 | 0.226307 |
| Connection Thal-R-PuM – Thal-L-VLp | T(51) = 2.39 | 0.020806 | 0.226655 |
| Connection Thal-R-CM – Thal-L-MDm | T(51) = -2.29 | 0.026431 | 0.234578 |
| Connection Thal-R-MDl – Thal-L-PuA | T(51) = 2.03 | 0.047567 | 0.285400 |
| Connection Thal-R-MDm – Thal-L-PuM | T(51) = 2.09 | 0.041289 | 0.292776 |
| Connection Thal-L-CM – Thal-R-VLp | T(51) = -2.10 | 0.040517 | 0.303161 |
| Connection DAN – Thal-L-LP | T(51) = -2.15 | 0.035935 | 0.323024 |
| Connection Thal-R-PuA – Thal-L-VLp | T(51) = 2.09 | 0.041459 | 0.359308 |
| Connection Visual – Thal-L-MDm | T(51) = -2.26 | 0.028175 | 0.366275 |
| Connection Thal-L-LGN – Thal-R-MDm | T(51) = -2.38 | 0.021124 | 0.422477 |
| Connection Thal-L-LGN – Thal-R-MDl | T(51) = -2.12 | 0.038489 | 0.422477 |
| Connection Thal-R-PuL – Thal-L-VLp | T(51) = 2.45 | 0.017716 | 0.441362 |
| Connection Thal-R-PuL – Thal-L-VLp | T(51) = 2.45 | 0.017716 | 0.441362 |
| Connection Visual – Thal-R-MDm | T(51) = -2.03 | 0.047457 | 0.462709 |
| Connection Thal-L-PuM – Thal-L-LP | T(51) = 2.34 | 0.023509 | 0.549938 |
| Connection Thal-L-PuM – Thal-L-MDm | T(51) = 2.19 | 0.033370 | 0.549938 |
| Connection Thal-L-PuM – Thal-L-VLp | T(51) = 2.11 | 0.039666 | 0.549938 |
| Connection Thal-L-PuA – Thal-L-MDl | T(51) = 2.47 | 0.016818 | 0.563525 |
| **Cluster 8/130** | **F(1,51) = 12.90** | **0.000738** | **0.011985** |
| Connection SMl – CON | T(51) = -3.59 | 0.000738 | 0.019176 |
| **Cluster 9/130** | **F(2,50) = 8.07** | **0.000920** | **0.013290** |
| Connection Thal-L-PuL – CON | T(51) = 2.83 | 0.006701 | 0.040208 |
| Connection GP-p-R – CON | T(51) = 3.56 | 0.000808 | 0.063005 |
| Connection Thal-R-PuA – CON | T(51) = 3.52 | 0.000911 | 0.071078 |
| Connection CON – HIP-body-L | T(51) = -2.96 | 0.004646 | 0.120797 |
| Connection GP-a-R – CON | T(51) = 2.26 | 0.027924 | 0.207863 |
| Connection CON – Thal-R-LGN | T(51) = -2.59 | 0.012406 | 0.217671 |
| Connection CON – HIP-body-R | T(51) = -2.52 | 0.014883 | 0.217671 |
| Connection CON – Thal-L-PuA | T(51) = 2.47 | 0.016744 | 0.217671 |
| Connection CON – DAN | T(51) = 2.24 | 0.029369 | 0.254529 |
| Connection PMN – CON | T(51) = 2.36 | 0.021949 | 0.285341 |
| Connection HIP-body-L – CON | T(51) = -2.03 | 0.047801 | 0.286806 |
| Connection DAN – CON | T(51) = 2.02 | 0.048767 | 0.335462 |
| Connection Thal-L-PuI – CON | T(51) = 2.03 | 0.047452 | 0.400962 |
| Connection GP-p-L – CON | T(51) = 2.02 | 0.048859 | 0.925414 |
| **Cluster 10/130** | **F(2,50) = 7.62** | **0.001297** | **0.016661** |
| Connection DMN – Thal-R-MDm | T(51) = -3.14 | 0.002788 | 0.022539 |
| Connection DMN – Thal-R-MDl | T(51) = -2.81 | 0.007086 | 0.035002 |
| Connection DMN – Thal-L-LP | T(51) = 2.71 | 0.009191 | 0.042172 |
| Connection Thal-L-LP – DMN | T(51) = 2.59 | 0.012554 | 0.089018 |
| Connection Thal-L-MDm – DMN | T(51) = 2.45 | 0.017765 | 0.159604 |
| Connection Thal-R-MDl – DMN | T(51) = -2.18 | 0.033734 | 0.219273 |
| Connection Thal-L-MDl – DMN | T(51) = 2.32 | 0.024327 | 0.557397 |
| **Cluster 11/130** | **F(2,50) = 7.36** | **0.001574** | **0.016661** |
| Connection DMN – HIP-head-l-L | T(51) = -4.21 | 0.000103 | 0.008053 |
| Connection DMN – HIP-head-l-R | T(51) = -3.86 | 0.000320 | 0.009876 |
| Connection DMN – AMY-l-LH | T(51) = -3.44 | 0.001174 | 0.015264 |
| Connection DMN – AMY-m-L | T(51) = -3.33 | 0.001617 | 0.015766 |
| Connection DMN – AMY-l-R | T(51) = -2.80 | 0.007180 | 0.035002 |
| Connection DMN – HIP-head-m1-L | T(51) = -2.59 | 0.012628 | 0.052874 |
| Connection DMN – AMY-m-R | T(51) = -2.49 | 0.016084 | 0.059740 |
| **Cluster 12/130** | **F(2,50) = 7.18** | **0.001813** | **0.016661** |
| Connection HIP-tail-R – HIP-head-m2-R | T(51) = 4.21 | 0.000103 | 0.004021 |
| Connection HIP-tail-L – HIP-head-m2-R | T(51) = 3.74 | 0.000474 | 0.020284 |
| Connection Thal-L-PuI – HIP-head-m2-R | T(51) = 3.14 | 0.002797 | 0.109092 |
| Connection HIP-head-m2-L – HIP-tail-L | T(51) = 3.33 | 0.001612 | 0.109826 |
| Connection HIP-head-m2-L – PMN | T(51) = -3.14 | 0.002816 | 0.109826 |
| Connection HIP-head-m2-L – HIP-tail-R | T(51) = 2.95 | 0.004815 | 0.125195 |
| Connection HIP-body-L – HIP-head-m2-R | T(51) = 2.49 | 0.016049 | 0.156478 |
| Connection HIP-head-m2-R – HIP-body-L | T(51) = 3.15 | 0.002717 | 0.201472 |
| Connection HIP-head-m2-R – HIP-tail-R | T(51) = 2.91 | 0.005334 | 0.201472 |
| Connection HIP-head-m2-R – HIP-body-R | T(51) = 2.77 | 0.007749 | 0.201472 |
| Connection Thal-L-CM – HIP-head-m2-L | T(51) = 2.31 | 0.024764 | 0.241448 |
| Connection HIP-body-R – HIP-head-m2-L | T(51) = 2.12 | 0.038764 | 0.251345 |
| Connection HIP-head-m2-L – Thal-R-CM | T(51) = 2.35 | 0.022936 | 0.277534 |
| Connection HIP-body-L – HIP-head-m2-L | T(51) = 2.06 | 0.044523 | 0.286806 |
| Connection HIP-head-m2-L – HIP-body-L | T(51) = 2.16 | 0.035407 | 0.298794 |
| Connection Thal-L-CM – HIP-head-m2-R | T(51) = 2.01 | 0.049791 | 0.303161 |
| Connection HIP-head-m2-L – HIP-body-R | T(51) = 2.04 | 0.046661 | 0.303297 |
| Connection HIP-head-m2-R – Thal-R-PuM | T(51) = -2.48 | 0.016647 | 0.324610 |
| Connection HIP-head-m2-R – HIP-tail-L | T(51) = 2.34 | 0.023204 | 0.361986 |
| Connection HIP-head-m2-R – Thal-L-PuM | T(51) = -2.12 | 0.038521 | 0.382523 |
| Connection HIP-head-m2-R – Thal-R-PuL | T(51) = -2.12 | 0.039233 | 0.382523 |
| **Cluster 13/130** | **F(2,50) = 7.17** | **0.001822** | **0.016661** |
| Connection CAU-tail-L – CON | T(51) = 3.56 | 0.000825 | 0.010722 |
| Connection CAU-body-R – CON | T(51) = 3.13 | 0.002887 | 0.028144 |
| Connection CAU-tail-R – CON | T(51) = 2.73 | 0.008749 | 0.341225 |
| **Cluster 14/130** | **F(2,50) = 7.12** | **0.001909** | **0.016661** |
| Connection Thal-R-CM – AMY-m-R | T(51) = 3.99 | 0.000214 | 0.016682 |
| Connection HIP-head-m1-L – HIP-body-L | T(51) = 3.54 | 0.000854 | 0.025362 |
| Connection HIP-head-m1-L – HIP-body-R | T(51) = 3.46 | 0.001090 | 0.025362 |
| Connection HIP-head-m1-L – GP-p-L | T(51) = 3.16 | 0.002657 | 0.036745 |
| Connection HIP-body-L – AMY-m-R | T(51) = 3.16 | 0.002651 | 0.058085 |
| Connection Thal-R-CM – AMY-l-R | T(51) = 3.36 | 0.001497 | 0.058370 |
| Connection Thal-R-CM – AMY-l-L | T(51) = 3.13 | 0.002896 | 0.060584 |
| Connection HIP-body-L – AMY-l-L | T(51) = 2.99 | 0.004318 | 0.067354 |
| Connection AMY-m-R – HIP-body-R | T(51) = 3.35 | 0.001516 | 0.067605 |
| Connection AMY-m-R – Thal-L-VPL | T(51) = -3.31 | 0.001733 | 0.067605 |
| Connection PMN – HIP-head-l-R | T(51) = -3.54 | 0.000877 | 0.068392 |
| Connection DAN – AMY-l-R | T(51) = 3.52 | 0.000924 | 0.072100 |
| Connection PMN – HIP-head-l-L | T(51) = -3.21 | 0.002270 | 0.088548 |
| Connection HIP-head-l-L – HIP-body-L | T(51) = 3.45 | 0.001143 | 0.089116 |
| Connection AMY-m-R – HIP-body-L | T(51) = 2.99 | 0.004270 | 0.091868 |
| Connection AMY-m-R – PMN | T(51) = -2.90 | 0.005444 | 0.091868 |
| Connection HIP-head-m1-L – GP-p-R | T(51) = 2.65 | 0.010580 | 0.096922 |
| Connection HIP-head-m1-L – Thal-L-CM | T(51) = 2.59 | 0.012426 | 0.096922 |
| Connection Thal-R-CM – AMY-m-L | T(51) = 2.74 | 0.008491 | 0.110378 |
| Connection AMY-l-L – GP-a-L | T(51) = -2.72 | 0.008970 | 0.112692 |
| Connection AMY-l-L – DAN | T(51) = 2.49 | 0.015892 | 0.112692 |
| Connection AMY-l-L – Thal-L-PuM | T(51) = -2.37 | 0.021798 | 0.130787 |
| Connection PMN – AMY-m-R | T(51) = -2.88 | 0.005739 | 0.135909 |
| Connection PMN – HIP-head-m1-L | T(51) = -2.73 | 0.008712 | 0.135909 |
| Connection Visual – HIP-head-m1-R | T(51) = 2.80 | 0.007194 | 0.140287 |
| Connection HIP-body-R – AMY-m-L | T(51) = 2.88 | 0.005857 | 0.144805 |
| Connection HIP-body-R – AMY-m-R | T(51) = 2.86 | 0.006179 | 0.144805 |
| Connection HIP-body-L – AMY-m-L | T(51) = 2.58 | 0.012667 | 0.156478 |
| Connection HIP-body-L – Thal-L-PuM | T(51) = -2.11 | 0.040189 | 0.156738 |
| Connection HIP-head-L – HIP-tail-L | T(51) = 2.07 | 0.043780 | 0.162610 |
| Connection AMY-l-L – Thal-R-CM | T(51) = 2.06 | 0.045017 | 0.167312 |
| Connection HIP-body-R – HIP-head-l-R | T(51) = 2.39 | 0.020334 | 0.176231 |
| Connection HIP-head-m1-R – HIP-tail-R | T(51) = 2.60 | 0.012151 | 0.200160 |
| Connection HIP-head-m1-R – HIP-body-R | T(51) = 2.56 | 0.013335 | 0.200160 |
| Connection HIP-head-m1-R – PON | T(51) = 2.54 | 0.014233 | 0.200160 |
| Connection AMY-m-R – Thal-R-PuA | T(51) = -2.12 | 0.039308 | 0.216094 |
| Connection Thal-L-CM – AMY-m-L | T(51) = 2.60 | 0.012135 | 0.216833 |
| Connection Thal-L-CM – HIP-head-m1-R | T(51) = 2.50 | 0.015734 | 0.216833 |
| Connection Thal-L-CM-SS – AMY-m-R | T(51) = 2.48 | 0.016679 | 0.216833 |
| Connection HIP-head-m1-R – HIP-body-L | T(51) = 2.35 | 0.022431 | 0.218705 |
| Connection HIP-head-l-L – Thal-R-CM | T(51) = 2.74 | 0.008543 | 0.222108 |
| Connection Thal-L-CM – HIP-head-m1-L | T(51) = 2.33 | 0.023902 | 0.241448 |
| Connection HIP-body-R – HIP-head-m1-R | T(51) = 2.11 | 0.039439 | 0.251345 |
| Connection Thal-R-PuL – HIP-head-l-L | T(51) = -3.08 | 0.003286 | 0.256325 |
| Connection Thal-R-LGN – AMY-l-R | T(51) = 2.22 | 0.030629 | 0.269010 |
| Connection Thal-R-LGN – AMY-l-L | T(51) = 2.10 | 0.041030 | 0.269010 |
| Connection HIP-tail-R – AMY-m-R | T(51) = 2.05 | 0.045974 | 0.275842 |
| Connection Thal-L-CM – AMY-l-R | T(51) = 2.08 | 0.042841 | 0.303161 |
| Connection DAN – HIP-head-m1-R | T(51) = 2.37 | 0.021360 | 0.323024 |
| Connection DAN – AMY-m-L | T(51) = 2.28 | 0.026818 | 0.323024 |
| Connection GP-p-R – AMY-l-L | T(51) = 2.37 | 0.021392 | 0.353543 |
| Connection HIP-head-l-L – HIP-body-R | T(51) = 2.20 | 0.032214 | 0.358960 |
| Connection AMY-l-R – Thal-L-PuM | T(51) = -2.61 | 0.011970 | 0.366128 |
| Connection AMY-l-R – DAN | T(51) = 2.23 | 0.030416 | 0.366128 |
| Connection AMY-l-R – GP-p-L | T(51) = 2.09 | 0.041650 | 0.366128 |
| Connection AMY-l-R – Thal-R-VPL | T(51) = -2.03 | 0.047332 | 0.366128 |
| Connection PON – HIP-head-l-R | T(51) = 2.16 | 0.035653 | 0.373809 |
| Connection PON – AMY-m-L | T(51) = 2.13 | 0.038169 | 0.373809 |
| Connection PON – AMY-m-R | T(51) = 2.07 | 0.043132 | 0.373809 |
| Connection Visual – AMY-l-R | T(51) = 2.18 | 0.033609 | 0.374497 |
| Connection HIP-head-l-L – PMN | T(51) = -2.06 | 0.044349 | 0.378178 |
| Connection Thal-L-LGN – AMY-l-R | T(51) = 2.69 | 0.009759 | 0.422477 |
| Connection Thal-R-PuL – HIP-head-m1-L | T(51) = -2.15 | 0.036318 | 0.441362 |
| Connection HIP-tail-L – HIP-head-l-R | T(51) = 2.07 | 0.043409 | 0.457540 |
| Connection AMY-m-L – PMN | T(51) = -2.39 | 0.020725 | 0.497065 |
| Connection AMY-m-L – DAN | T(51) = 2.36 | 0.021909 | 0.497065 |
| Connection AMY-m-L – GP-a-L | T(51) = -2.13 | 0.038026 | 0.497065 |
| Connection AMY-m-L – Thal-L-VPL | T(51) = -2.06 | 0.044390 | 0.497065 |
| Connection HIP-head-l-R – DAN | T(51) = 2.74 | 0.008484 | 0.499825 |
| Connection HIP-head-l-R – PON | T(51) = 2.40 | 0.019930 | 0.499825 |
| **Cluster 15/130** | **F(2,50) = 7.11** | **0.001922** | **0.016661** |
| Connection DMN – HIP-body-R | T(51) = -3.81 | 0.000380 | 0.009876 |
| Connection DMN – HIP-tail-L | T(51) = -3.60 | 0.000712 | 0.013883 |
| Connection DMN – Thal-R-Pul | T(51) = -3.45 | 0.001127 | 0.015264 |
| Connection DMN – Thal-R-PuM | T(51) = -3.36 | 0.001482 | 0.015766 |
| Connection DMN – Visual | T(51) = -3.13 | 0.002890 | 0.022539 |
| Connection DMN – HIP-tail-R | T(51) = -2.95 | 0.004762 | 0.031368 |
| Connection DMN – HIP-body-L | T(51) = -2.93 | 0.005070 | 0.031368 |
| Connection DMN – PMN | T(51) = -2.85 | 0.006222 | 0.034663 |
| Connection DMN – Thal-R-CM-SS-8206 | T(51) = -2.58 | 0.012880 | 0.052874 |
| Connection PMN – DMN | T(51) = -2.78 | 0.007555 | 0.135909 |
| Connection Thal-R-CM – DMN | T(51) = -2.29 | 0.026098 | 0.234578 |
| Connection DAN – DMN | T(51) = -2.11 | 0.040124 | 0.323024 |
| Connection Thal-R-PuA – DMN | T(51) = -2.24 | 0.029819 | 0.344350 |
| Connection Thal-R-PuL – DMN | T(51) = -2.04 | 0.046561 | 0.441362 |
| Connection Thal-L-PuM – DMN | T(51) = 2.43 | 0.018569 | 0.549938 |
| **Cluster 16/130** | **F(2,50) = 6.44** | **0.003248** | **0.026394** |
| Connection FPN – Thal-L-VA | T(51) = -2.94 | 0.004893 | 0.051191 |
| Connection FPN – PUT-VP-R | T(51) = 2.88 | 0.005825 | 0.051191 |
| Connection FPN – Thal-R-VA | T(51) = -2.85 | 0.006280 | 0.051191 |
| Connection PUT-VP-R – FPN | T(51) = 2.86 | 0.006112 | 0.059595 |
| Connection FPN – Thal-R-VLa | T(51) = -2.68 | 0.009880 | 0.060242 |
| Connection FPN – PUT-DP-R | T(51) = 2.52 | 0.014874 | 0.077343 |
| **Cluster 17/130** | **F(2,50) = 5.90** | **0.004991** | **0.037757** |
| Connection FPN – Thal-R-MDl | T(51) = -4.23 | 0.000099 | 0.003852 |
| Connection FPN – Thal-R-VLp | T(51) = -3.50 | 0.000982 | 0.019143 |
| Connection FPN – Thal-R-MDm | T(51) = -3.15 | 0.002716 | 0.037592 |
| Connection FPN – Thal-L-MDl | T(51) = -2.45 | 0.017674 | 0.081577 |
| Connection FPN – Thal-L-MDm | T(51) = -2.38 | 0.020858 | 0.081577 |
| Connection FPN – Thal-L-VLa | T(51) = -2.28 | 0.027079 | 0.093877 |
| Connection Thal-L-LP – FPN | T(51) = -2.46 | 0.017529 | 0.105174 |
| **Cluster 18/130** | **F(1,51) = 8.52** | **0.005228** | **0.037757** |
| Connection DMN – CON | T(51) = -2.92 | 0.005228 | 0.031368 |
| Connection CON – DMN | T(51) = -3.05 | 0.003640 | 0.120797 |
| **Cluster 19/130** | **F(2,50) = 5.65** | **0.006120** | **0.041875** |
| Connection Thal-L-CeM – SAN | T(51) = 2.23 | 0.030425 | 0.469049 |
| **Cluster 20/130** | **F(2,50) = 5.40** | **0.007535** | **0.047821** |
| Connection CAU-body-L – CAU-tail-R | T(51) = 2.69 | 0.009531 | 0.082606 |
| Connection CAU-body-L – CAU-body-R | T(51) = 2.59 | 0.012505 | 0.097538 |
| Connection CAU-VA-L – CAU-tail-L | T(51) = 2.86 | 0.006046 | 0.127264 |
| Connection CAU-VA-L – CAU-tail-R | T(51) = 2.72 | 0.008953 | 0.127264 |
| Connection CAU-VA-R – CAU-tail-L | T(51) = 2.65 | 0.010713 | 0.233022 |
| Connection CAU-VA-R – CAU-tail-R | T(51) = 2.52 | 0.014937 | 0.233022 |
| **Cluster 21/130** | **F(2,50) = 5.37** | **0.007725** | **0.047821** |
| Connection CAU-VA-L – SAN | T(51) = 4.09 | 0.000152 | 0.011822 |
| Connection CAU-body-L – SAN | T(51) = 2.39 | 0.020504 | 0.145395 |
| Connection CAU-VA-R – SAN | T(51) = 2.09 | 0.041257 | 0.321075 |
| **Cluster 22/130** | **F(2,50) = 5.28** | **0.008340** | **0.049284** |
| Connection FPN – CAU-body-R | T(51) = -3.68 | 0.000564 | 0.014662 |
| Connection FPN – CAU-tail-L | T(51) = -2.32 | 0.024528 | 0.091103 |
| Connection CAU-tail-L – FPN | T(51) = -2.35 | 0.022704 | 0.160994 |

**Supplementary Table 2.** Detailed results for the group-level functional connectivity analysis obtained using a ROI-level p-FDR correction (ROI mass/intensity) false-positive control method implemented in the CONN toolbox, showing mass. The table lists significant results for ROI pairs with a *p*-uncorrected connection threshold of 0.05 and a cluster-level p-FDR corrected threshold of 0.05.

| **Analysis Unit** | **Score** | **Statistic** | **p-unc** | **p-FDR** | **p-FWE** |
| --- | --- | --- | --- | --- | --- |
| **ROI 1/38 Thal-L-PuL**  Mass = 169.69  Size = 14 | 52.63 |  | 0.000000  0.000039  0.000921 | 0.000000  0.001488  0.017500 | 0.000000  0.001000  0.031000 |
| Connection Thal-L-PuL – Thal-R-VLp | 52.63 | T(51) = 4.53 | 0.000036 | 0.020154 |  |
| Connection Thal-L-PuL – Thal-R-PuA |  | T(51) = 4.36 | 0.000063 | 0.020154 |  |
| Connection Thal-L-PuL – Thal-L-MDl |  | T(51) = 4.34 | 0.000068 | 0.020154 |  |
| Connection Thal-L-PuL – Thal-R-PuM |  | T(51) = 3.72 | 0.000500 | 0.042250 |  |
| Connection Thal-L-PuL – Thal-L-MDm |  | T(51) = 3.71 | 0.000507 | 0.042250 |  |
| Connection Thal-L-PuL – Thal-L-VLp |  | T(51) = 3.60 | 0.000720 | 0.044006 |  |
| Connection Thal-L-PuL – Thal-R-MDl |  | T(51) = 3.58 | 0.000755 | 0.044218 |  |
| Connection Thal-L-PuL – Thal-R-MDm |  | T(51) = 3.40 | 0.001299 | 0.065210 |  |
| Connection Thal-L-PuL – Thal-L-PuM |  | T(51) = 3.23 | 0.002167 | 0.087049 |  |
| Connection Thal-L-PuL – Thal-R-VPL |  | T(51) = 3.19 | 0.002448 | 0.095591 |  |
| Connection Thal-L-PuL – Thal-R-PuL |  | T(51) = 2.92 | 0.005240 | 0.117656 |  |
| Connection Thal-L-PuL – Thal-L-PuA |  | T(51) = 2.66 | 0.010539 | 0.176399 |  |
| Connection Thal-L-PuL – Thal-R-VLa |  | T(51) = 2.31 | 0.024770 | 0.263833 |  |
| Connection Thal-L-PuL – Thal-L-VPL |  | T(51) = 2.18 | 0.033589 | 0.301342 |  |
| **ROI 2/38 Thal-L-LP**  Mass = 122.20  Size = 13 | 34.64 |  | 0.000289  0.000260  0.001526 | 0.005500  0.004946  0.019333 | 0.010000  0.009000  0.050000 |
| Connection Thal-L-LP – Thal-L-MDm |  | T(51) = 4.32 | 0.000072 | 0.020154 |  |
| Connection Thal-L-LP – Thal-L-VLp |  | T(51) = 4.02 | 0.000195 | 0.030420 |  |
| Connection Thal-L-LP – Thal-L-MDl |  | T(51) = 3.62 | 0.000675 | 0.044006 |  |
| Connection Thal-L-LP – Thal-L-VLa |  | T(51) = 3.44 | 0.001163 | 0.062884 |  |
| Connection Thal-L-LP – Thal-R-VLp |  | T(51) = 3.38 | 0.001412 | 0.068463 |  |
| Connection Thal-L-LP – Thal-L-CeM |  | T(51) = 3.01 | 0.004055 | 0.106215 |  |
| Connection Thal-L-LP – Thal-L-AV |  | T(51) = 2.88 | 0.005764 | 0.119417 |  |
| Connection Thal-L-LP – Thal-R-MDm |  | T(51) = 2.88 | 0.005775 | 0.119417 |  |
| Connection Thal-L-LP – Thal-L-VA |  | T(51) = 2.53 | 0.014518 | 0.210431 |  |
| Connection Thal-L-LP – Thal-L-PuA |  | T(51) = 2.30 | 0.025820 | 0.269917 |  |
| Connection Thal-L-LP – Thal-R-PuM |  | T(51) = 2.20 | 0.032659 | 0.301342 |  |
| Connection Thal-L-LP – VPL |  | T(51) = 2.18 | 0.033642 | 0.301342 |  |
| Connection Thal-L-LP – Thal-R-LGN |  | T(51) = -2.03 | 0.048114 | 0.330778 |  |
| **ROI 3/38 PUT-VA-L**  Mass = 107.97  Size = 14 | 26.71 |  | 0.001789  0.000658  0.000921 | 0.011333  0.008333  0.017500 | 0.058000  0.022000  0.031000 |
| Connection PUT-VA-L – Thal-R-VLp |  | T(51) = 3.63 | 0.000649 | 0.044006 |  |
| Connection PUT-VA-L – PUT-DA-L |  | T(51) = 3.61 | 0.000689 | 0.044006 |  |
| Connection PUT-VA-L – Thal-L-MDm |  | T(51) = 3.09 | 0.003217 | 0.100906 |  |
| Connection PUT-VA-L – PUT-VP-R |  | T(51) = 2.99 | 0.004301 | 0.106798 |  |
| Connection PUT-VA-L – PUT-DA-R |  | T(51) = 2.98 | 0.004406 | 0.106798 |  |
| Connection PUT-VA-L – Thal-R-LP |  | T(51) = 2.90 | 0.005456 | 0.119417 |  |
| Connection PUT-VA-L – Thal-R-MDl |  | T(51) = 2.80 | 0.007144 | 0.140061 |  |
| Connection PUT-VA-L – Thal-L-VLp |  | T(51) = 2.49 | 0.016260 | 0.215442 |  |
| Connection PUT-VA-L – PUT-VP-L |  | T(51) = 2.40 | 0.020284 | 0.241694 |  |
| Connection PUT-VA-L – PUT-DP-R |  | T(51) = 2.34 | 0.023221 | 0.259873 |  |
| Connection PUT-VA-L – PUT-DP-L |  | T(51) = 2.33 | 0.023658 | 0.259873 |  |
| Connection PUT-VA-L – Thal-L-AV |  | T(51) = 2.28 | 0.026871 | 0.273770 |  |
| Connection PUT-VA-L – Thal-L-MDl |  | T(51) = 2.26 | 0.028139 | 0.280593 |  |
| Connection PUT-VA-L – Thal-R-MDm |  | T(51) = 2.22 | 0.031194 | 0.292396 |  |
| **ROI 4/38 Thal-R-LP**  Mass = 93.84  Size = 11 | 27.55 |  | 0.001500  0.001400  0.004263 | 0.011333  0.013300  0.032400 | 0.054000  0.045000  0.122000 |
| Connection Thal-R-LP – Thal-R-VLp |  | T(51) = 4.21 | 0.000103 | 0.024081 |  |
| Connection Thal-R-LP – Thal-L-VLp |  | T(51) = 3.79 | 0.000401 | 0.042250 |  |
| Connection Thal-R-LP – Thal-L-MDl |  | T(51) = 3.15 | 0.002755 | 0.096835 |  |
| Connection Thal-R-LP – Thal-R-VLa |  | T(51) = 3.14 | 0.002826 | 0.096918 |  |
| Connection Thal-R-LP – Thal-R-MDl |  | T(51) = 3.12 | 0.002950 | 0.098756 |  |
| Connection Thal-R-LP – Thal-L-VPL |  | T(51) = 2.51 | 0.015183 | 0.211271 |  |
| Connection Thal-R-LP – Thal-R-MDm |  | T(51) = 2.43 | 0.018498 | 0.226163 |  |
| Connection Thal-R-LP – Thal-L-VLa |  | T(51) = 2.38 | 0.021070 | 0.243446 |  |
| Connection Thal-R-LP – Thal-L-MDm |  | T(51) = 2.26 | 0.027829 | 0.279479 |  |
| Connection Thal-R-LP – Thal-R-PuA |  | T(51) = 2.24 | 0.029566 | 0.284654 |  |
| Connection Thal-R-LP – Thal-L-PuA |  | T(51) = 2.05 | 0.045649 | 0.330778 |  |
| **ROI 5/38 Thal-R-PuM**  Mass = 80.41  Size = 9 | 26.71 |  | 0.001789  0.003026  0.010842 | 0.011333  0.023000  0.068667 | 0.058000  0.088000  0.279000 |
| Connection Thal-R-PuM – Thal-L-LP |  | T(51) = 4.56 | 0.000032 | 0.020154 |  |
| Connection Thal-R-PuM – Thal-L-PuL |  | T(51) = 3.92 | 0.000267 | 0.034152 |  |
| Connection Thal-R-PuM – Thal-L-VPL |  | T(51) = 3.17 | 0.002543 | 0.096636 |  |
| Connection Thal-R-PuM – Thal-R-VA |  | T(51) = -2.76 | 0.007970 | 0.151325 |  |
| Connection Thal-R-PuM – Thal-R-LGN |  | T(51) = 2.75 | 0.008226 | 0.152189 |  |
| Connection Thal-R-PuM – Thal-L-VLp |  | T(51) = 2.39 | 0.020806 | 0.243446 |  |
| Connection Thal-R-PuM – Thal-R-PuL |  | T(51) = 2.18 | 0.034028 | 0.302807 |  |
| Connection Thal-R-PuM – Thal-L-PuA |  | T(51) = 2.09 | 0.041646 | 0.321723 |  |
| Connection Thal-R-PuM – Thal-L-VA |  | T(51) = -2.05 | 0.045947 | 0.330778 |  |
| **ROI 6/38 PUT-DP-R**  Mass = 76.38  Size = 8 | 27.81 |  | 0.001447  0.003822  0.017763 | 0.011333  0.024208  0.084375 | 0.051000  0.113000  0.409000 |
| Connection PUT-DP-R – PUT-DA-L |  | T(51) = 4.11 | 0.000145 | 0.025796 |  |
| Connection PUT-DP-R – PUT-DA-R |  | T(51) = 4.10 | 0.000147 | 0.025796 |  |
| Connection PUT-DP-R – PUT-VP-L |  | T(51) = 3.49 | 0.001010 | 0.056817 |  |
| Connection PUT-DP-R – Thal-R-PuA |  | T(51) = -2.73 | 0.008709 | 0.156979 |  |
| Connection PUT-DP-R – Thal-L-AV |  | T(51) = 2.68 | 0.009853 | 0.168938 |  |
| Connection PUT-DP-R – PUT-VP-R |  | T(51) = 2.54 | 0.014031 | 0.205503 |  |
| Connection PUT-DP-R – Thal-R-CM |  | T(51) = -2.29 | 0.026229 | 0.271158 |  |
| Connection PUT-DP-R – Thal-R-AV |  | T(51) = 2.04 | 0.046797 | 0.330778 |  |
| **ROI 7/38 Thal-R-MDm**  Mass = 71.47  Size = 11 | 18.31 |  | 0.014895  0.005289  0.004263 | 0.066250  0.028714  0.032400 | 0.377000  0.153000  0.122000 |
| Connection Thal-R-MDm – Thal-R-VLp |  | T(51) = 3.67 | 0.000577 | 0.043939 |  |
| Connection Thal-R-MDm – Thal-R-AV |  | T(51) = 3.15 | 0.002722 | 0.096835 |  |
| Connection Thal-R-MDm – Thal-L-VLp |  | T(51) = 2.57 | 0.013128 | 0.200630 |  |
| Connection Thal-R-MDm – Thal-R-MDl |  | T(51) = 2.52 | 0.014924 | 0.211271 |  |
| Connection Thal-R-MDm – Thal-L-PuL |  | T(51) = 2.46 | 0.017147 | 0.221177 |  |
| Connection Thal-R-MDm – Thal-R-VLa |  | T(51) = 2.37 | 0.021854 | 0.249815 |  |
| Connection Thal-R-MDm – Thal-L-MDm |  | T(51) = 2.32 | 0.024662 | 0.263833 |  |
| Connection Thal-R-MDm – Thal-L-AV |  | T(51) = 2.25 | 0.028577 | 0.282956 |  |
| Connection Thal-R-MDm – Thal-L-MDl |  | T(51) = 2.12 | 0.039050 | 0.321723 |  |
| Connection Thal-R-MDm – Thal-L-PuM |  | T(51) = 2.09 | 0.041289 | 0.321723 |  |
| Connection Thal-R-MDm – Thal-R-LP |  | T(51) = 2.03 | 0.047420 | 0.330778 |  |
